# *E. coli* TraR allosterically regulates transcription initiation by altering RNA polymerase conformation and dynamics

**DOI:** 10.1101/766725

**Authors:** James Chen, Saumya Gopalkrishnan, Courtney Chiu, Albert Y. Chen, Elizabeth A. Campbell, Richard L. Gourse, Wilma Ross, Seth A. Darst

## Abstract

TraR and its homolog DksA are bacterial proteins that regulate transcription initiation by binding directly to RNA polymerase (RNAP) rather than to promoter DNA. Effects of TraR mimic the combined effects of DksA and its cofactor ppGpp. How TraR and its homologs regulate transcription is unclear. Here, we use cryo-electron microscopy to determine structures of *Escherichia coli* RNAP, with or without TraR, and of an RNAP-promoter complex. TraR binding induced RNAP conformational changes not seen in previous crystallographic analyses, and a quantitative analysis of RNAP conformational heterogeneity revealed TraR-induced changes in RNAP dynamics. These changes involve mobile regions of RNAP affecting promoter DNA interactions, including the βlobe, the clamp, the bridge helix, and several lineage-specific insertions. Using mutational approaches, we show that these structural changes, as well as effects on σ^70^ region 1.1, are critical for transcription activation or inhibition, depending on the kinetic features of regulated promoters.

## Introduction

Transcription initiation is a major control point for gene expression. In bacteria, a single catalytically active RNA polymerase (RNAP) performs all transcription, but a σ factor is required for promoter utilization (Burgess et al., 1969; Feklistov et al., 2014). In *Escherichia coli* (*Eco*), the essential primary σ factor, σ^70^, binds to RNAP to form the σ^70^-holoenzyme (Eσ^70^) that is capable of recognizing and initiating at promoters for most genes. Upon locating the promoter, Eσ^70^ melts a ∼13 bp segment of DNA to form the open promoter complex (RPo) in which the DNA template strand (t-strand) is loaded into the RNAP active site, exposing the transcription start site (Bae et al., 2015b; Zuo and Steitz, 2015). A key feature of the RPo formation pathway is that it is a multi-step process, with the RNAP-promoter complex passing through multiple intermediates before the final, transcription competent RPo is formed (Hubin et al., 2017a; Ruff et al., 2015; Saecker et al., 2002).

A variety of transcription factors bind to the promoter DNA and/or to RNAP directly to regulate initiation (Browning and Busby, 2016; Haugen et al., 2008). Bacterial RNAP-binding factors, encoded by the chromosome or by bacteriophage or extrachromosomal elements, interact with several different regions of the enzyme to regulate its functions (Haugen et al., 2008). One such factor is ppGpp, a modified nucleotide that functions together with the RNAP-binding protein DksA in *Eco* to reprogram bacterial metabolism in response to nutritional stresses during the so-called stringent response. Following amino acid starvation, ppGpp is synthesized by the RelA factor in response to uncharged tRNAs in the ribosomal A site (Brown et al., 2016; Cashel and Gallant, 1969; Ryals et al., 1982). Together, ppGpp and DksA alter the expression of as many as 750 genes within 5 minutes of ppGpp induction (Paul et al., 2004a; 2005; Sanchez-Vazquez et al., 2019), inhibiting, for example, promoters responsible for ribosome biogenesis and activating promoters responsible for amino acid synthesis.

The overall RNAP structure is reminiscent of a crab claw, with one pincer comprising primarily the β’ subunit, and the other primarily the β subunit (Zhang et al., 1999). Between the two pincers is a large cleft that contains the active site. In Eσ^70^ without nucleic acids, this channel is occupied by the σ^70^ domain which is ejected upon entry of the downstream duplex DNA (Bae et al., 2013; Mekler et al., 2002). The Bridge Helix (BH) bridges the two pincers across the cleft, separating the cleft into the main channel, where σ^70^ or nucleic acids reside, and the secondary channel, where NTPs enter the RNAP active site.

DksA binds in the RNAP secondary channel (Lennon et al., 2012; Molodtsov et al., 2018; Perederina et al., 2004). ppGpp binds directly to RNAP at two binding sites: site 1, located at the interface of the β′ and ω subunits (Ross et al., 2013; Zuo et al., 2013), and site 2, located at the interface of β′ and DksA (Molodtsov et al., 2018; Ross et al., 2016). The ppGpp bound at site 1 inhibits transcription ∼2-fold under conditions where the effects of ppGpp bound at both sites together with DksA are as much as 20-fold (Paul et al., 2004b; Ross et al., 2016). By contrast, DksA and ppGpp bound at site 2 are necessary and sufficient for activation (Ross et al., 2016).

TraR is a distant homolog of DksA. Although only half the length of DksA, TraR regulates *Eco* transcription by binding to the RNAP secondary channel and mimicking the combined effects of DksA and ppGpp (Blankschien et al., 2009; Gopalkrishnan et al., 2017). TraR is encoded by the conjugative F plasmid and is expressed from the pY promoter as part of the major *tra* operon transcript (Frost et al., 1994; Maneewannakul and Ippen-Ihler, 1993). Like DksA, TraR inhibits Eσ^70^-dependent transcription from ribosomal RNA promoters (e.g. *rrnB* P1) and ribosomal protein promoters (e.g. *rpsT* P2, expressing S20), and activates amino acid biosynthesis and transport promoters (e.g. *pthrABC, phisG, pargI, plivJ*) *in vivo* and *in vitro* (Blankschien et al., 2009; Gopalkrishnan et al., 2017). The affinity of TraR for RNAP is only slightly higher than that of DksA, yet its effects on promoters negatively regulated by DksA/ppGpp *in vitro* are as large or larger than those of DksA/ppGpp (Gopalkrishnan et al., 2017). The effects of TraR on promoters positively regulated by ppGpp/DksA are also independent of ppGpp (Gopalkrishnan et al., 2017).

Models for DksA/ppGpp and TraR binding to RNAP have been proposed based on biochemical and genetic approaches (Gopalkrishnan et al., 2017; Parshin et al., 2015; Ross et al., 2013; 2016). Crystal structures of DksA/ppGpp/RNAP and TraR/RNAP confirmed the general features of these models and provided additional detail about their interactions with RNAP, but did not reveal the mechanism of inhibition or activation, in large part because of crystal packing constraints on the movement of mobile regions of the complex (Molodtsov et al., 2018). Thus, the structural basis for the effects of DksA/ppGpp or TraR on transcription have remained elusive.

To help understand TraR regulation and principles of the regulation of transcription initiation in general, we used single particle cryo-electron microscopy (cryo-EM) to examine structures of Eσ^70^ alone, Eσ^70^ bound to TraR (TraR-Eσ^70^), and Eσ^70^ bound to a promoter inhibited by TraR [*rpsT* P2; (Gopalkrishnan et al., 2017)]. Cryo-EM allows the visualization of multiple conformational states populated in solution and in the absence of crystal packing constraints. Furthermore, new software tools allow for the analysis of molecular motions in the cryo-EM data (Nakane et al., 2018).

The TraR-Eσ^70^ structures show TraR binding in the secondary channel of the RNAP, consistent with the TraR-Eσ^70^ model (Gopalkrishnan et al., 2017) and crystal structure (Molodtsov et al., 2018). However, the cryo-EM structures reveal major TraR-induced changes to the RNAP conformation that were not evident in the crystal structure due to crystal packing constraints. Structural analyses generated mechanistic hypotheses for TraR function in both activation and inhibition of transcription that were then tested biochemically. On the basis of the combined structural and functional analyses, we propose a model in which TraR accelerates multiple steps along the RPo formation pathway while at the same time modulates the relative stability of intermediates in the pathway. Whether a promoter is activated or inhibited by TraR is determined by the intrinsic kinetic properties of the promoter (Galburt, 2018; Haugen et al., 2008; Paul et al., 2005).

## Results

### Cryo-EM structures of TraR-Eσ^70^

We used single-particle cryo-EM to examine *Eco* TraR-Eσ^70^ in the absence of crystal packing interactions that could influence the conformational states. TraR function in cryo-EM solution conditions (Chen et al., 2019) was indistinguishable from standard *in vitro* assay conditions (Figure 1 - figure supplement 1A-C). Analysis of the cryo-EM data gave rise to three distinct conformational classes (Figure 1 - figure supplement 1D). All three structures are essentially identical except for the disposition of Si3 [also called β’i6; (Lane and Darst, 2010a)], a 188-residue lineage-specific insertion (LSI) in the trigger-loop (TL) of *Eco* RNAP (Chlenov et al., 2005) (Figures 1A, B). The first class [TraR-Eσ^70^(I)] contained approximately 41% of the particles and resolved to a nominal resolution of 3.7 Å (Figure 1A). The second class [TraR-Eσ^70^(II)] contained approximately 33% of the particles and resolved to a nominal resolution of 3.8 Å (Figure 1B). The third class [TraR-Eσ^70^(III)] contained the remaining 26% of the particles and resolved to a nominal resolution of 3.9 Å (Figure 1 - figure supplement 1D, Figure 1 - figure supplement 2; Supplementary file 1). With Si3 (β’ residues 948-1126) excluded, the structures superimpose with a root-mean-square deviation (rmsd) of 0.495 Å over 3,654 α-carbons.

**Figure 1.**
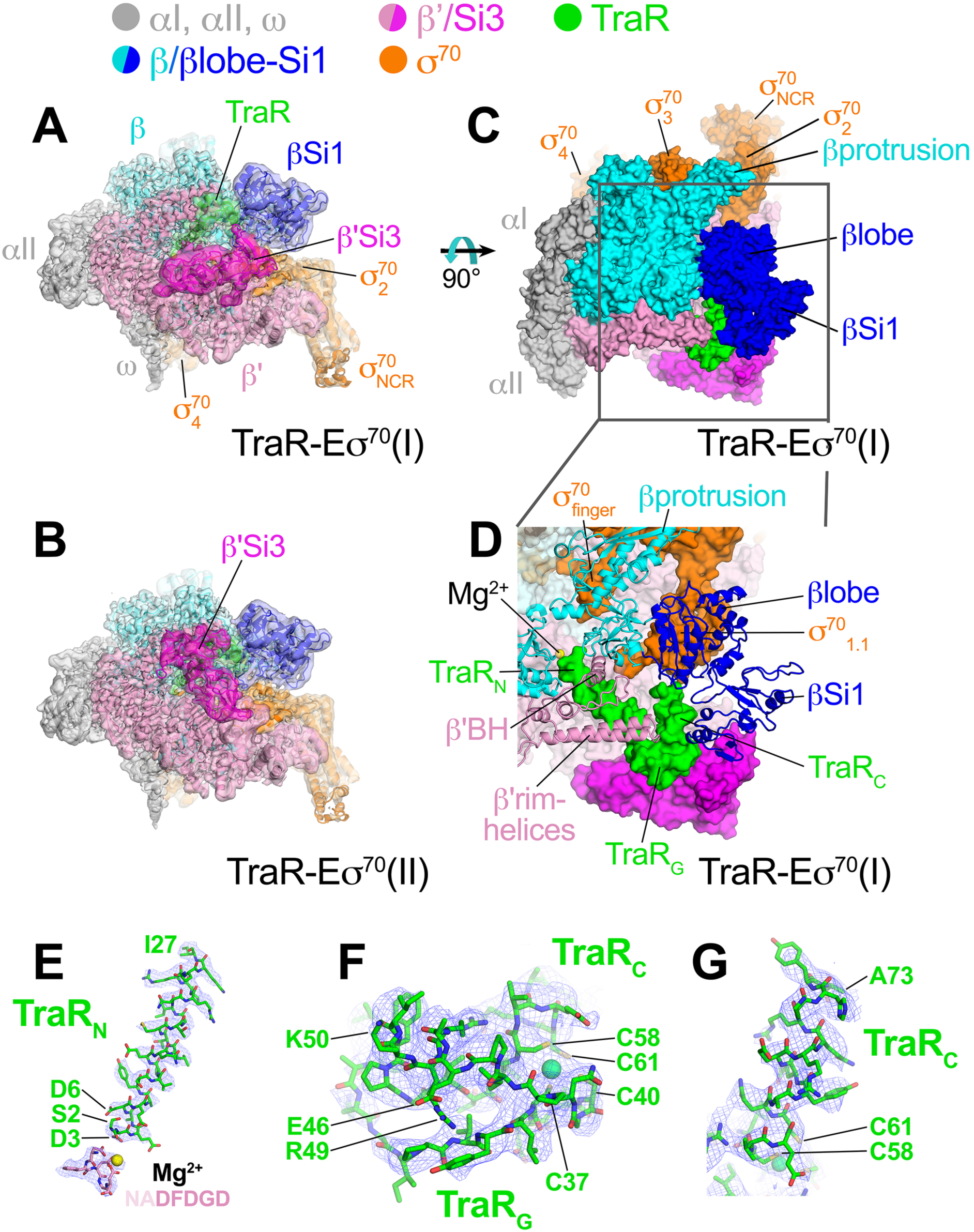
Cryo-EM structure of TraR-Eσ^70^. (*top*) Color-coding key. (**A**) TraR-Eσ^70^(I) - cryo-EM density map (3.7 Å nominal resolution, low-pass filtered to the local resolution) is shown as a transparent surface and colored according to the key. The final model is superimposed. (**B**) TraR-Eσ^70^(II) - cryo-EM density map (3.8 Å nominal resolution, low-pass filtered to the local resolution) is shown as a transparent surface and colored according to the key. The final model is superimposed. (**C**) Top view of TraR-Eσ^70^(I). The boxed area is magnified in (**D**). (**D**) Magnified top view of TraR-Eσ^70^(I) - shows TraR_N_ (starting near RNAP active site Mg^2+^, extending out secondary channel), TraR_G_ (interacting primarily with β’rim-helices), and TraRT_C_ (interacting with βlobe-Si1). (**E - G**) Cryo-EM density (blue mesh) defining the TraR structure. (**E**) TraR_N_ and -NADFDGD- motif of RNAP β’ (chelating active site Mg^2+^). (**F**) TraR_G_. (**G**) TraR _C_.

The overall binding mode of TraR in the cryo-EM structures (Figures 1A-D) is consistent with the effects of TraR or RNAP substitutions on TraR function (Gopalkrishnan et al., 2017) and is broadly consistent with the X-ray structure (Molodtsov et al., 2018). TraR can be divided into three structural elements, an N-terminal helix (TraR_N_, residues 2-27; Figures 1D, E), a globular domain that includes a 4-Cys Zn^2+^-binding motif (TraR_G_, residues 28-57; Figures 1D, F), and a C-terminal helix (TraR_C_, residues 58-73; Figures 1D, G). TraR_N_ extends from the RNAP active site out through the RNAP secondary channel to the β’rim-helices (at the entrance to the RNAP secondary channel), interacting with key RNAP structural elements surrounding the active site, including the -NADFDGD-motif that chelates the active site Mg^2+^ (Zhang et al., 1999), the F-loop (Miropolskaya et al., 2009), and the bridge-helix (Figure 1D). The N-terminal tip of TraR_N_ (TraR residue S2) is only 4.3 Å from the active site Mg^2+^ (Figure 1E). TraR_G_ interacts primarily with the β’rim-helices at the entrance of the secondary channel (Figure 1D).

The interactions of TraR_C_ with RNAP differ between the cryo-EM and X-ray structures due to conformational changes induced by TraR binding detected by the cryo-EM structure that were not observed in the X-ray structure (see below). Indeed, the cryo-EM and X-ray structures superimpose with an rmsd of 4.26 Å over 3,471 α-carbons, indicating significant conformational differences.

### Cryo-EM analysis of Eσ^70^ and *rpsT* P2 RPo

To understand how TraR-induced conformational changes regulate RPo formation, we analysed single-particle cryo-EM data for Eσ^70^ alone and Eσ^70^ bound to the *rpsT* P2 promoter. Our aim was to explore conformational space and dynamics unencumbered by crystal packing constraints to compare with the TraR-Eσ^70^ data above. Cryo-EM data for Eσ^70^ resolved to a nominal resolution of 4.1 Å (Figure 1 - figure supplements 3, 4; Supplementary file 1).

Analysis of the *rpsT* P2-RPo cryo-EM data gave rise to two conformational classes that differed only in the disposition of the upstream promoter DNA and αCTDs (Figure 2 - figure supplement 1). We focus here on the highest resolution class at a nominal resolution of 3.4 Å (Figure 2B, Figure 2 - figure supplement 1, Figure 2 - figure supplement 2; Supplementary file 1).

**Figure 2.**
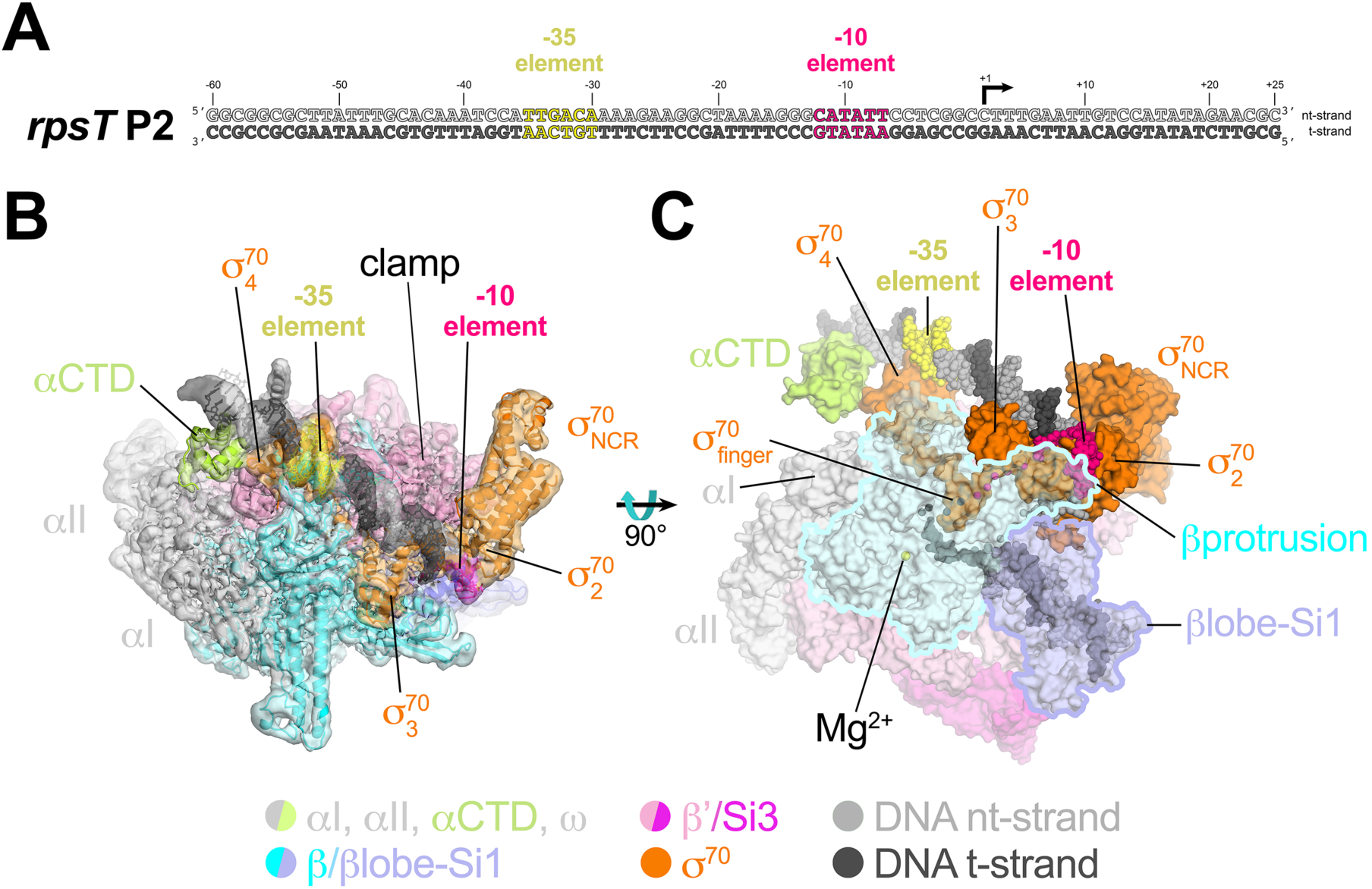
Cryo-EM structure of *rpsT* P2-RPo. (**A**) The *Eco rpsT* P2 promoter fragment used for cryo-EM. (**B**) *rpsT* P2-RPo cryo-EM density map (3.4 Å nominal resolution, low-pass filtered to the local resolution) is shown as a transparent surface and colored according to the key. The final model is superimposed. The DNA was modeled from -45 to +21. The t-strand DNA from -10 to -2, and the nt-strand DNA from -3 to +2 were disordered. (**C**) Top view of *rpsT* P2-RPo. DNA is shown as atomic spheres. Proteins are shown as molecular surfaces. Much of the β subunit is transparent to reveal the active site Mg^2+^ (yellow sphere), σ^70^_finger_, and DNA inside the RNAP active site cleft.

The closed-clamp RNAP in the *rpsT* P2-RPo interacts with the promoter DNA in the same way as seen in other RPo structures determined by X-ray crystallography (Bae et al., 2015b; 2015a; Hubin et al., 2017b) or cryo-EM (Boyaci et al., 2019) and is consistent with the DNase I footprint of the *rpsT* P2 RPo (Gopalkrishnan et al., 2017). In the *rpsT* P2-RPo structure we observed an α-subunit C-terminal domain [αCTD; (Ross et al., 1993)] bound to the promoter DNA minor groove (Benoff et al., 2002; Ross et al., 2001) just upstream of the promoter -35 element [-38 to -43, corresponding to the proximal UP element subsite (Estrem et al., 1999)]. This αCTD interacts with σ^70^ through an interface previously characterized through genetic analyses (Ross et al., 2003) (Figures 2B, C). The αCTDs are linked to the α-N-terminal domains (αNTDs) by ∼15-residue flexible linkers (Blatter et al., 1994; Jeon et al., 1995). Density for the residues connecting the αCTD and αNTD was not observed in the cryo-EM map.

Comparing the RNAP conformations of the TraR-Eσ^70^, Eσ^70^, and *rpsT* P2-RPo cryo-EM structures revealed key differences that suggest how TraR activates and inhibits transcription. Below we outline these differences and present experients that test their implications for function.

### β’Si3 is in two conformations, one of which is important for TraR activation function

The three TraR-Eσ^70^ structures differ from each other only in the disposition of Si3. Si3 comprises two tandem repeats of the sandwich-barrel hybrid motif (SBHM) fold (Chlenov et al., 2005; Iyer et al., 2003), SBHMa and SBHMb (Figure 3A). Si3 is linked to the TL-helices by extended, flexible linkers. In TraR-Eσ^70^(I) and TraR-Eσ^70^(II), Si3 is in two distinct positions with respect to the RNAP (Figures 1A, 1B, 3A), while in TraR-Eσ^70^(III) Si3 is disordered (Figure 1 - figure supplement 1D). Si3 in the TraR-Eσ^70^(I) structure [Si3(I)] interacts primarily with the β’shelf (SBHMa) and the β’jaw (SBHMb) in a manner seen in many previous *Eco* RNAP X-ray (Bae et al., 2013) and cryo-EM structures (Chen et al., 2017; Kang et al., 2017; Liu et al., 2017). Si3 in the TraR-Eσ^70^(II) structure [Si3(II)] is rotated 121° such that SBHMa interacts with the β’jaw and SBHMb interacts with TraR_G_ (Figures 3A-C), a disposition of Si3 that, to our knowledge, has not been observed previously.

**Figure 3.**
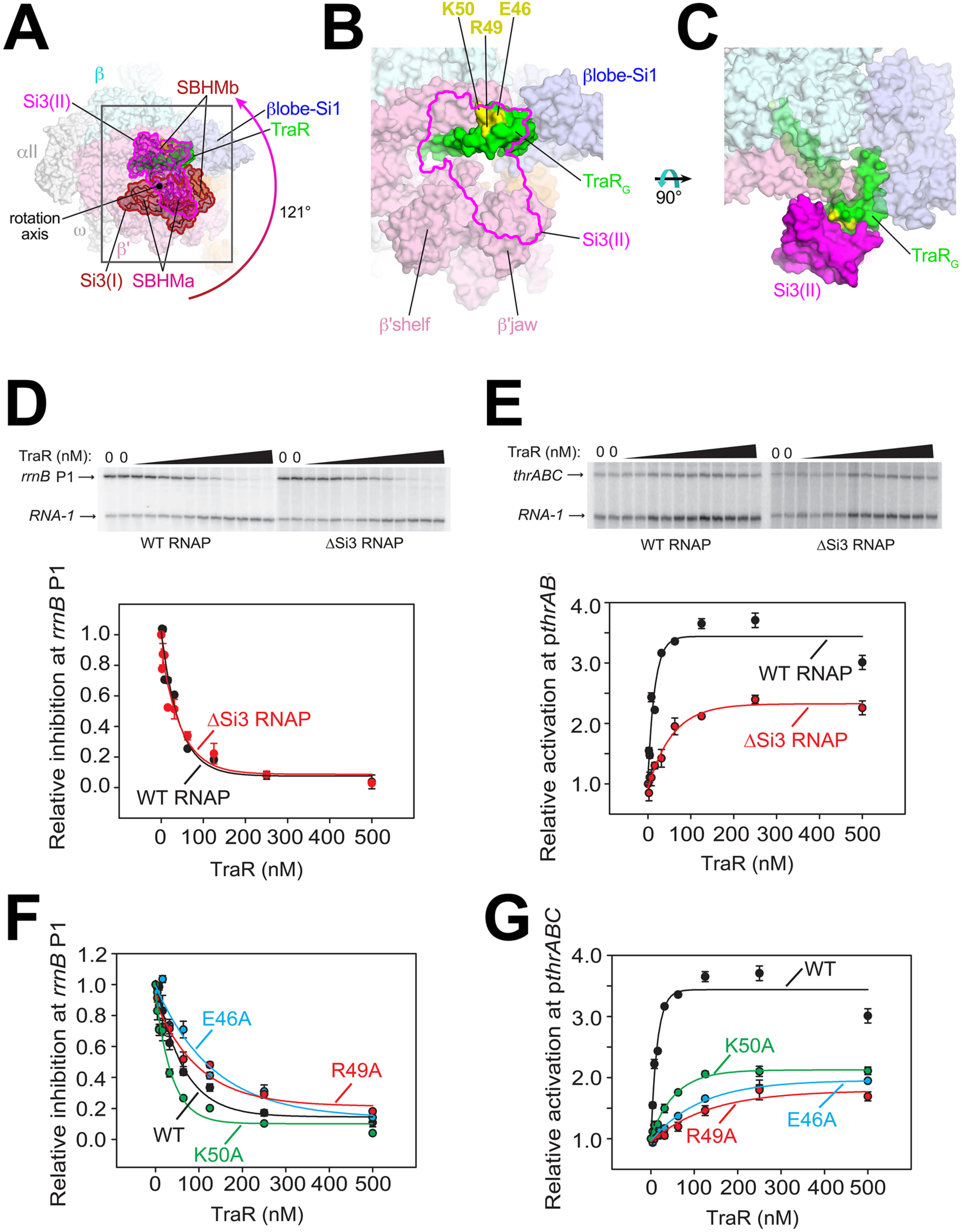
Conformational flexibility of β’Si3 in TraR-Eσ^70^. (**A**) Overall view of TraR-Eσ^70^ structure with alternative positions of Si3. Si3(I) is shown in brown. A ∼121° rotation about the rotation axis shown gives rise to the position of Si3(II) shown in magenta. Si3 comprises two SBHM domains (Chlenov et al., 2005; Iyer et al., 2003), denoted SBHMa and SBHMb. The boxed region is magnified in (**B**). (**B**) Magnified view of TraR-Eσ^70^(II) [same view as (**A**)]. The position of Si3(II) is outlined in magenta but the rest of Si3 is removed, revealing TraR behind. Three residues central to the TraR-Si3(II) interface (TraR-E46, R49, and K50) are colored yellow. (**C**) Orthogonal view as (**B**), showing the extensive TraR-Si3(II) interface. (**D**) – (**G**): Si3 interaction with TraR_G_ affects activation but not inhibition. Quantifications show averages with range from two independent experiments. (**D**) (top) Multi round *in vitro* transcription of *rrnB* P1 over a range of TraR concentrations (wedge indicates 2 nM - 2 µM) in the presence of WT-RNAP or ΔSi3-RNAP as indicated. Plasmid templates also contained the RNA-1 promoter. (bottom) Quantification of transcripts from experiments like those shown on (top) plotted relative to values in the absence of TraR. The IC_50_ for inhibition by TraR was ∼40 nM for both data sets. (**E**) (top) Multi round *in vitro* transcription of *thrABC* over a range of TraR concentrations (wedge indicates 2 nM - 2 µM) in the presence of 20 nM WT-RNAP or ΔSi3-RNAP as indicated. Plasmid templates also contained the RNA-1 promoter. (bottom) Quantification of transcripts from experiments like those shown on (top) plotted relative to values in the absence of TraR. (**F**) and (**G**) Multi round *in vitro* transcription of *rrnB* P1 (**F**) or p*thrABC* (**G**) was performed with 20 nM WT-Eσ^70^ at a range of concentrations of WT or variant TraR (2 nM -2 µM). Transcripts were quantified and plotted relative to values in the absence of any factor (n=2). For (**F**), IC_50_ for inhibition by WT-TraR was ∼50 nM, by E46A TraR was ∼115 nM, R49A TraR was ∼85 nM and by K50A TraR was ∼30 nM.

To test if this alternative conformation [Si3(II)] is relevant to TraR function, we compared TraR-mediated function at promoters known to be inhibited or activated by TraR with wild-type (WT) and ΔSi3-RNAP. Deletion of Si3 had little to no effect on TraR-mediated inhibition of *rrnB* P1 and *rpsT* P2 (Figure 3D, Figure 3 - figure supplement 1A) but transcription by ΔSi3-RNAP was activated only ∼50% compared with WT-RNAP on three different TraR-activated promoters (p*thrABC*, Figure 3E; p*argI*, Figure 3 - figure supplement 1B; p*hisG*, Figure 3 - figure supplement 1C).

Three TraR_G_ residues (TraR-E46, R49, and K50) are central to the Si3-TraR_G_ interface (Figure 3B, C). Individual alanine substitutions of these TraR residues (TraR-E46A, R49A, or K50A) gave rise to similar results as deleting Si3. Inhibition of *rrnB* P1 was similar to WT-TraR for TraR-K50A, and mildly impaired for TraR-E46A or R49A (Figure 3F; legend for IC_50_ values). Maximal inhibition was achieved at higher E46A or R49A TraR concentrations. However, these same variants exhibited at least ∼2-fold reduced activation at the *thrABC* promoter (Figure 3G) even at saturating TraR concentrations, indicating a role for the TraR-Si3 interaction in the mechanism of activation. Consistent with these results, these TraR variants were proficient in RNAP binding in a competition assay (Figure 3 - figure supplement 1F). By contrast, substitutions for nearby TraR variants P43A and P45A were defective for binding to RNAP, and their functional defects were overcome at higher TraR concentrations (Figure 3 - figure supplement 1D-F).

The combination of the TraR-Si3 interface mutants with the ΔSi3-RNAP was epistatic, i.e. no additional defects were observed over the ∼2-fold reduction in activation observed for the Si3-TraR interface mutants or the ΔSi3-RNAP individually (Figure 3 - figure supplement 1G). These results indicate that the Si3(SBHMb)-TraR_G_ interaction enabled by the Si3(II) conformation accounts for part of the TraR-mediated effect on activation.

### A TraR-induced ∼18° rotation of βlobe-Si1 plays a major role in transcription regulation

The large cleft between the two pincers in the struture of RNAP forms a channel that accommodates the downstream duplex DNA between the β’shelf and the clamp on one side, and the βlobe-Si1 domains on the other (Figure 2B, C). In Eσ^70^ without nucleic acids, this channel is occupied by the σ^70^ domain, which is ejected upon entry of the downstream duplex DNA (Figure 1D) (Bae et al., 2013; Mekler et al., 2002). TraR binding induces a ∼18° rotation of the RNAP βlobe-Si1 domains (the two domains move together as a rigid body), shifting the βlobe-Si1 towards TraR, allowing the βlobe-Si1 to establish an interface with TraR_G_ and TraR_C_ (615 Å^2^ interface area; Figure 4A).

**Figure 4.**
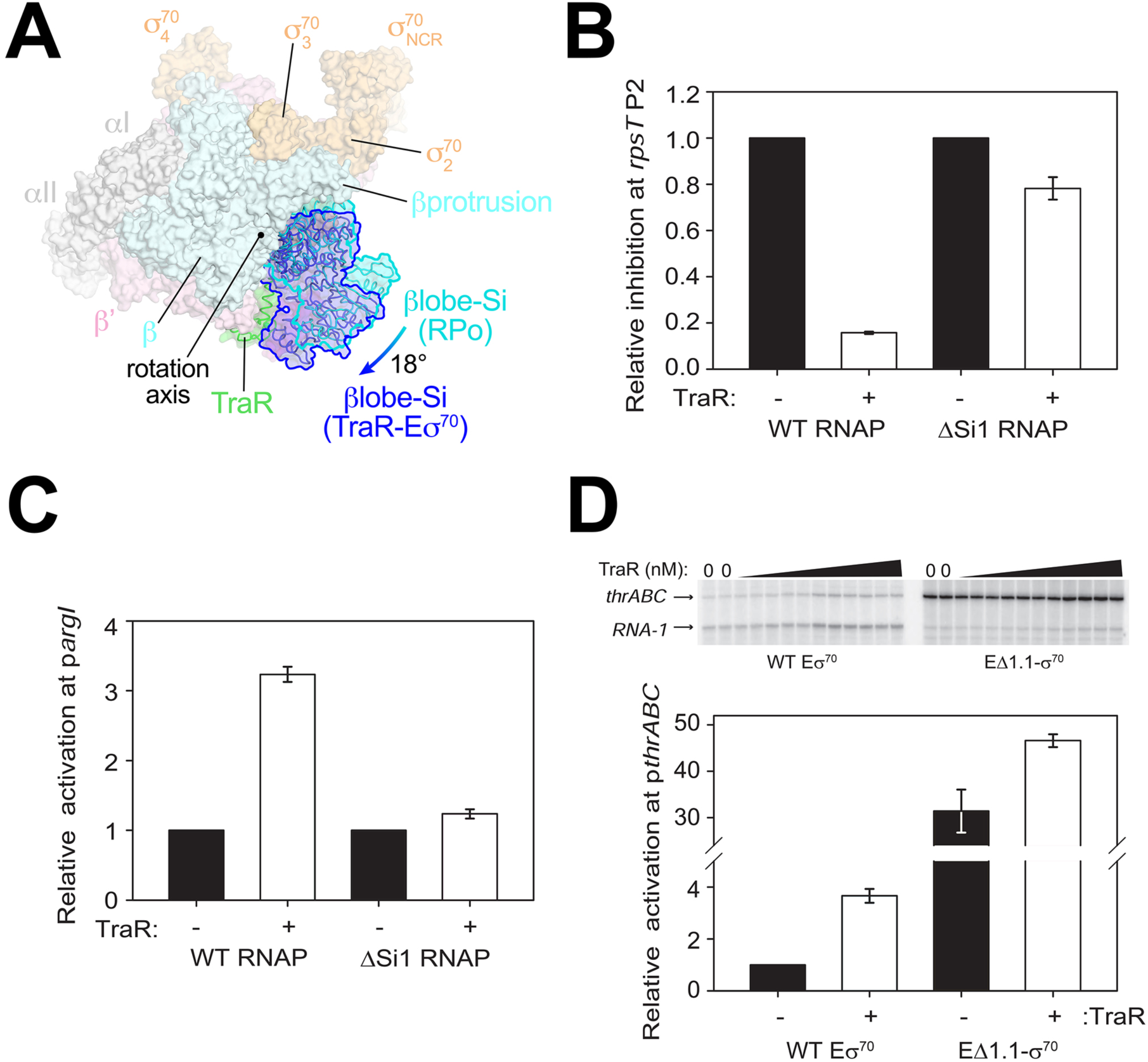
TraR and the βlobe-Si1 domain. (**A**) Overall top view of the TraR-Eσ^70^ structure with the βlobe-Si1 in dark blue. The corresponding position of the βlobe-Si1 in the *rpsT* P2-RPo structure (Figure 3) is shown in light blue. The βlobe-Si1 of the *rpsT* P2-RPo structure (light blue) undergoes an ∼19° rotation about the rotation axis shown to the βlobe-Si1 position in the TraR-Eσ^70^ structure (dark blue), generating an extensive TraR-βlobe-Si1 interface. (**B**) Transcription of inhibited promoter *rpsT* P2 by 20 nM WT-RNAP or ΔSi1-RNAP with (+) or without (-) 250 nM TraR as indicated. Error bars denote standard deviation of three independent measurements. (**C**) Transcription of activated promoter p*argI* by 20 nM WT-RNAP or ΔSi1-RNAP with (+) or without (-) 250 nM TraR as indicated. Error bars denote standard deviation of three independent measurements. (**D**) (top) Multi-round *in vitro* transcription was carried out at a range of TraR concentrations (wedge indicates 4 nM - 4 μM) in the presence of 20 nM WT-Eσ^70^ or EΔ1.1σ^70^ as indicated. Plasmid template also contained the RNA-1 promoter. (bottom) Transcripts from experiments such as those in (top) were quantified and plotted relative to values in the absence of TraR with WT-Eσ^70^ or EΔ1.1σ^70^ with (+) or without (-) 250 nM TraR as indicated. Averages with range from two independent experiments are shown.

Si1 [also called βi4; (Lane and Darst, 2010a)] is an LSI within the βlobe. Most of the TraR βlobe-Si1 interface (77%) is between TraR and Si1. Deleting Si1 from RNAP nearly abolishes activation function [p*argI,* Figure 4C; *thrABC*, (Gopalkrishnan et al., 2017)], even at saturating concentrations of TraR to overcome weakened TraR binding (Gopalkrishnan et al., 2017). These results suggest that the βlobe-Si1 rotation induced by TraR is essential to TraR-mediated activation.

The rotation of the βlobe-Si1 widens the gap between the βprotrusion and the βlobe (Figure 4A) and changes the shape of the RNAP channel, altering RNAP contacts with σ^70^1.1 in Eσ^70^. We hypothesize that altering the RNAP contacts with σ^70^ in the channel facilitates σ^70^ ejection during RPo formation, potentially contributing to activation of promoters that are limited at this step. To test this hypothesis, we investigated TraR function on an inhibited (*rrnB* P1) and an activated (*thrABC*) promoter with holoenzyme lacking σ^70^ (EΔ1.1σ^70^).

Eσ^70^ exhibited a low basal level of transcription from the TraR-activated *thrABC* promoter in the absence of TraR, and transcription was stimulated about ∼4-fold by TraR (Figure 4D). EΔ1.1σ^70^ exhibited a striking increase in basal transcription activity (∼32-fold) compared to WT-Eσ^70^ in the absence of TraR (Figure 4D). A small further increase in transcription was observed upon the addition of TraR (Figure 4D). These results suggest that σ^70^ is an obstacle to promoter DNA entering the RNAP channel and that TraR partially overcomes this barrier. In contrast to deletion of region σ^70^1.1, which bypasses the requirement for TraR, rotation of the βlobe-Si1 does not weaken σ^70^1.1-RNAP contacts sufficiently to release σ^70^1.1 completely (Figure 1D). Rather, βlobe-Si1 rotation facilitates the competition between promoter DNA and σ^70^ during RPo formation. Our results suggest that TraR-activated promoters are defined, in part, by being limited at the σ^70^ ejection step.

βSi1 was also required for inhibition of *rpsT* P2 (Figure 4B) and *rrnB* P1 transcription by TraR (Gopalkrishnan et al., 2017). However, in contrast to the effect of σ^70^1.1 on activation, deletion if σ^70^ had little effect on basal transcription from the TraR-inhibited *rrnB* P1 promoter, and inhibition of *rrnB* P1 with EΔ1.1σ^70^ by TraR was only slightly defective (Figure 4 - figure supplement 1A). Thus, in contrast to the effects of Si1 on activation by TraR, we suggest that the effect of TraR on inhibition of transcription involves the βlobe-Si1 domains but this is not mediated by σ^70^ (see Discussion). We propose that TraR-mediated stimulation of σ^70^ release still occurs at inhibited promoters like *rrnB* P1 and *rpsT* P2, but this has little effect on transcription because these pormoters are limited by their unstable RPo (Barker et al., 2001) (Figure 4-figure supplement 1).

### TraR induces β’shelf rotation and a bridge-helix kink, contributing to inhibition

TraR binding induces a ∼4.5° rotation of the β’shelf module (Figure 5A, B). The BH leads directly into the shelf module, and a kink is introduced in the BH, a long α-helix that traverses the RNAP active site cleft from one pincer to the other, directly across from the active site Mg^2+^ (Figure 5B, C). The BH plays critical roles in the RNAP nucleotide addition cycle (Lane and Darst, 2010b), including interacting with the t-strand DNA at the active site (Figure 5D). TraR causes the BH to kink towards the t-strand DNA (Figure 5C), similar to BH kinks observed previously (Tagami et al., 2011; 2010; Weixlbaumer et al., 2013; Zhang et al., 1999), resulting in a steric clash with the normal position of the t-strand nucleotide at +2 (Figure 5E). Thus, the TraR-induced BH kink would sterically prevent the proper positioning of the t-strand DNA in RPo, likely contributing to inhibition of transcription.

**Figure 5.**
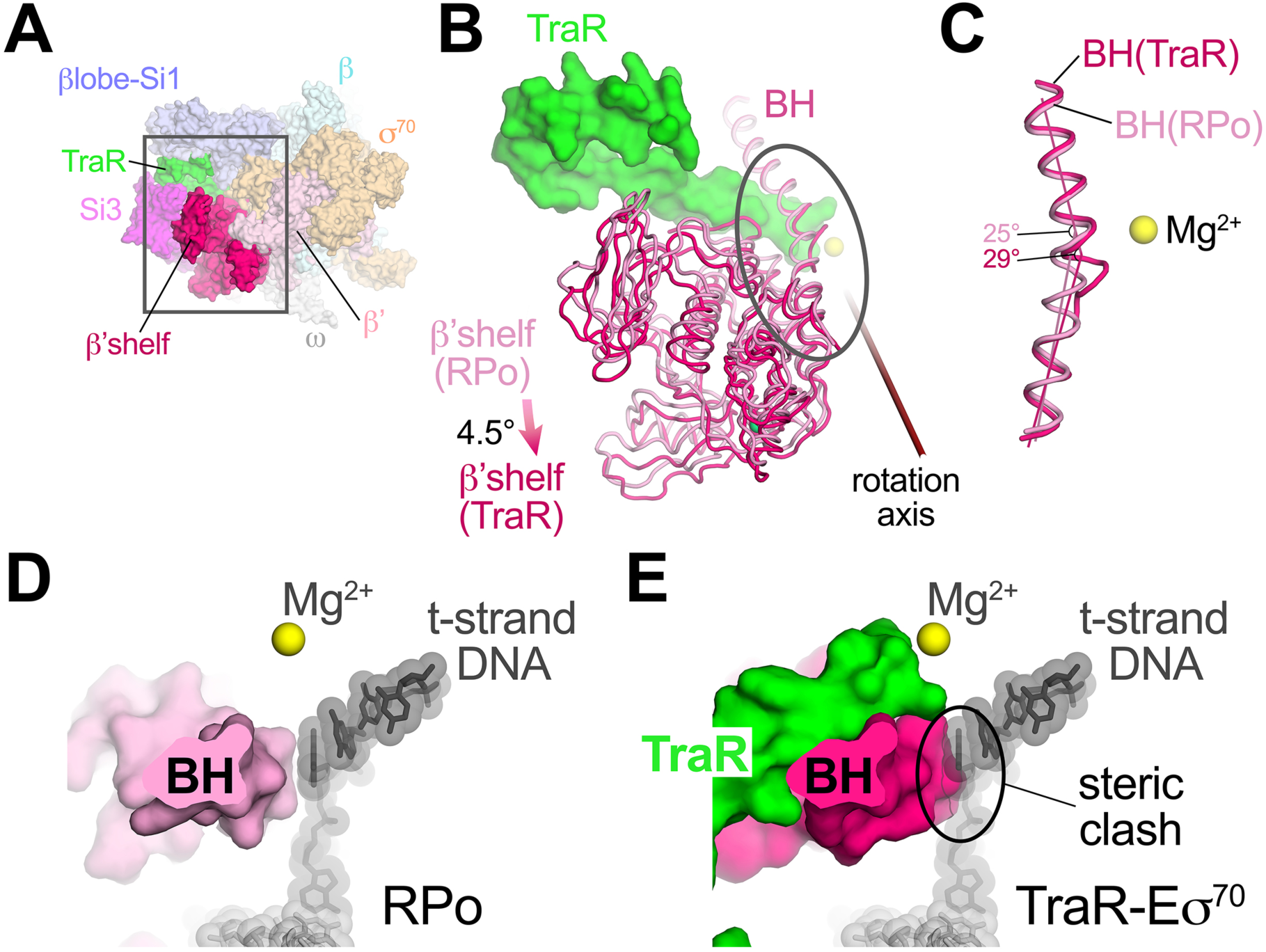
TraR rotates the β’shelf and kinks the BH. (**A**) Overall view if the TraR-Eσ^70^(I) structure, shown as a molecular surface. The β’shelf domain is highlighted in hot pink. The β’shelf (which here includes the β’jaw) comprises Eco β’ residues 787-931/1135-1150/1216-1317. The boxed region is magnified in (**B**). (**B**) Comparison of the *rpsT* P2-RPo BH-β’shelf (pink) and the TraR-Eσ^70^ BH-β’shelf (hot pink). Binding of TraR induces an ∼4.5° rotation (about the rotation axis shown) of the RPo-β’shelf to the position of the TraR-Eσ^70^ β’shelf and a kink in the BH (circled region, which is magnified in (**C**)). (**C**) Focus on the region of the BH kink, which is centered near β’L788. The kink in the RPo BH is about 25°, while the kink in the TraR-Eσ^70^ BH is about 29°. (**D**) View down the axis of the *rpsT* P2-RPo BH. The t-strand DNA, positioned at the RNAP active site (marked by the Mg^2+^ ion), closely approaches the BH. (**E**) View down the axis of the TraR-Eσ^70^ BH. The BH kink induced by TraR binding sterically clashes with the position of the t-strand DNA (superimposed from the RPo structure).

### TraR binding alters clamp dynamics, restricting clamp motions compared to Eσ^70^, likely stimulating transcription bubble nucleation

TraR induces conformational changes in the RNAP βlobe-Si1 (Figure 4A), β’shelf, and BH (Figure 5) structural modules. Although we noted modest changes in clamp positions (Supplementary file 2), we suspected that conformational heterogeneity of Eσ^70^ (limiting the resolution of the single particle analysis; Figure 1 - figure supplement 3) likely arose primarily from clamp motions that led to a continuous distribution of clamp positions that could not be easily classified into distinct conformational states, and that these motions were dampened in the TraR-Eσ^70^ and *rpsT* P2-RPo structures. We therefore analysed and compared the heterogeneity of RNAP clamp positions between the Eσ^70^, TraR-Eσ^70^, and *rpsT* P2-RPo datasets using multibody refinement as implemented in RELION 3 (Nakane et al., 2018). The maps used for multi-body refinement were carefully chosen to be equivalently processed. After initial classification to remove junk particles, particles were 3D auto-refined, then the refinement metadata and post-processing were used as inputs for RELION CTF refinement and Bayesian Polishing (Zivanov et al., 2018). After a final round of 3D auto-refinement (but no further classification), the *rpsT* P2-RPo dataset had the smallest number of particles (370,965), so a random subset of particles from the other datasets (TraR-Eσ^70^ and Eσ^70^) were processed so that each map for multi-body refinement was generated from the same number of particles (370,965). The final maps used for multi-body refinement had nominal resolutions of 4.0 Å (TraR-Eσ^70^; red dashed box in Figure 1 - figure supplement 1), 4.6 Å (Eσ^70^; red dashed box in Figure 1 - figure supplement 3), and 3.5 Å (*rpsT* P2-RPo; red dashed box in Figure 2 - figure supplement 1).

For Eσ^70^, three major components (Eigenvectors) of clamp motion were revealed (Figure 6A-D). For each Eigenvector, the histogram of Eigenvalues closely approximated a Gaussian distribution (Figure 6B-D). To quantitate the range of clamp motion represented by the Eigenvalues, we divided the particles into three bins according to their Eigenvalues such that each bin contained an equal number of particles (red, gray, and blue in Figure 6B-D). Three-dimensional alignments and reconstructions were then calculated for each bin.

**Figure 6.**
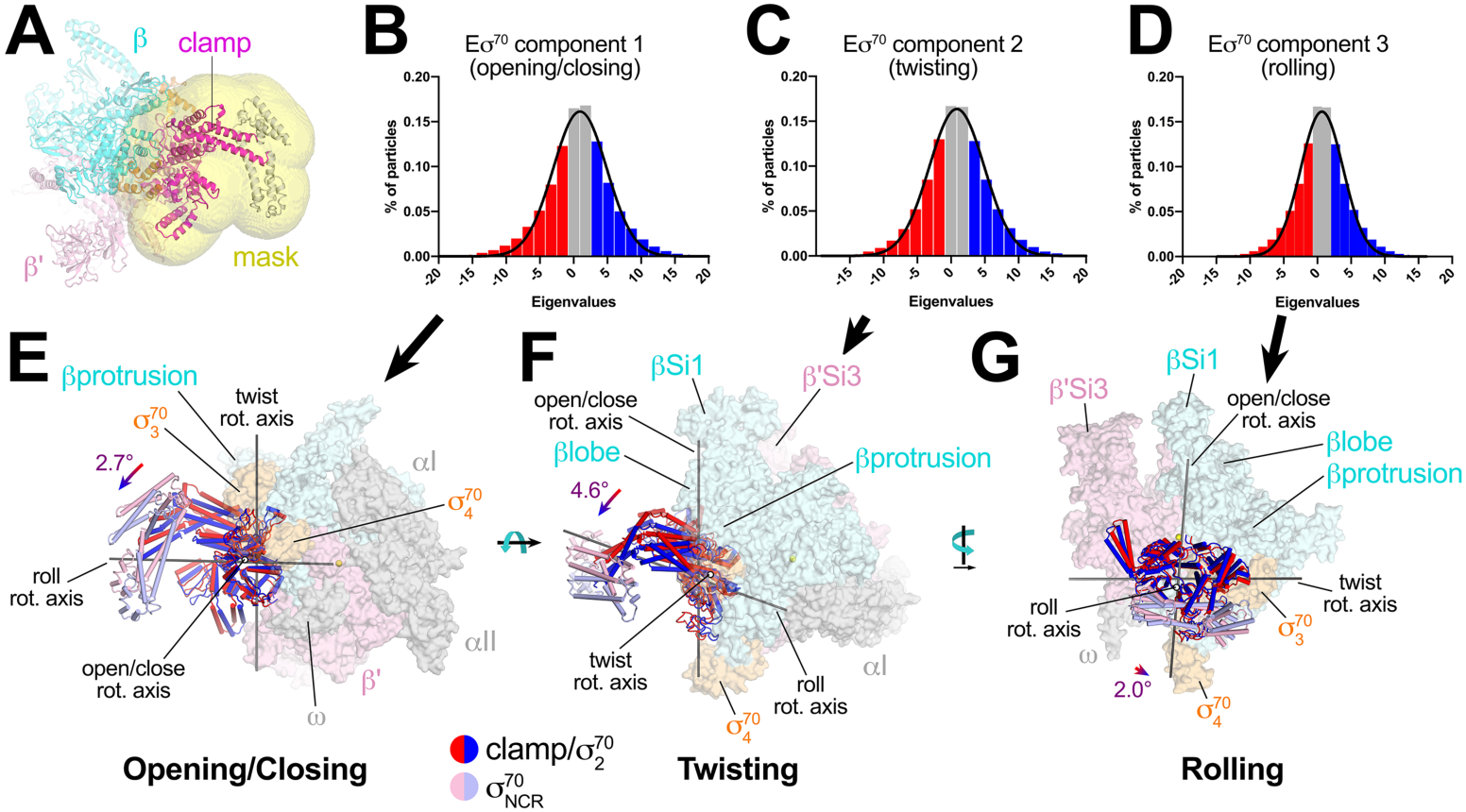
Multi-body analysis of Eσ^70^ clamp motions. **(A)** Model of Eσ^70^ refined into the consensus cryo-EM map (nominal 4.1 Å resolution). The RNAP clamp is highlighted in magenta. The clamp (which in the context of Eσ^70^ includes σ^70^2) comprises the following *Eco* RNAP residues: β1319-1342; β’ 1-342, 1318-1344; σ^70^ 92-137, 353-449. The mask used to analyze clamp motions by multi-body refinement (Nakane et al., 2018) is shown as a transparent yellow surface. **(B - D)** Histograms of Eigenvalue distributions (% of particles assigned each Eigenvalue from the dataset) for each of the three major principle components (Eigenvectors) from the multi-body analysis (Nakane et al., 2018). Each set of particles were divided into three equal-sized bins (Eigenvalue ≤ -2, red; -1 ≤ Eigenvalue ≤ 1, gray; Eigenvalue ≥ 2, blue). The solid lines denote Gaussian fits to the histograms. **(B)** Component 1. **(C)** Component 2. **(D)** Component 3. **(E - G)** Three-dimensional reconstructions were calculated from the red and blue-binned particles for each principle component and models were generated by rigid body refinement. The models were superimposed using α-carbons of the RNAP structural core, revealing the alternate clamp positions shown (red and blue α-carbon ribbons with cylindrical helices). The σ^70^NCR, attached to the clamp but not included in the clamp motion analyses, is shown in faded colors. For each component, the clamp rotation and the direction of the rotation axis were determined (rotation axes are shown in gray). **(E)** Component 1 - clamp opening/closing. **(F)** Component 2 - clamp twisting. **(G)** Component 3 - clamp rolling.

For component 1, the red and blue particles gave rise to reconstructions that differed in clamp positions by a rotation angle of 2.7° in a motion we call opening/closing (Figure 6E). The low Eigenvalue particles yielded a closed clamp (red), while the high Eigenvalue particles (blue) gave an open clamp. In the middle, the particles having intermediate Eigenvalues (gray) gave a clamp position half-way in between the red and the blue, as expected (not shown).

Component 2 gave rise to clamp positions that differed by a 4.6° rotation about a rotation axis roughly perpendicular to the open/close rotation axis, a motion we call twisting (Figure 6F). Finally, component 3 gave rise to clamp positions that differed by a 2.0° rotation about a third rotation axis parallel with the long axis of the clamp, a motion we call rolling (Figure 6G).

Using the parameters of the Gaussian fits to the Eigenvalue histograms (Figures 6B-D), we could estimate the full range of clamp rotations for each component, which we defined as the rotation range that accounted for 98% of the particles (excluding 1% of the particles at each tail; Figure 7).

**Figure 7.**
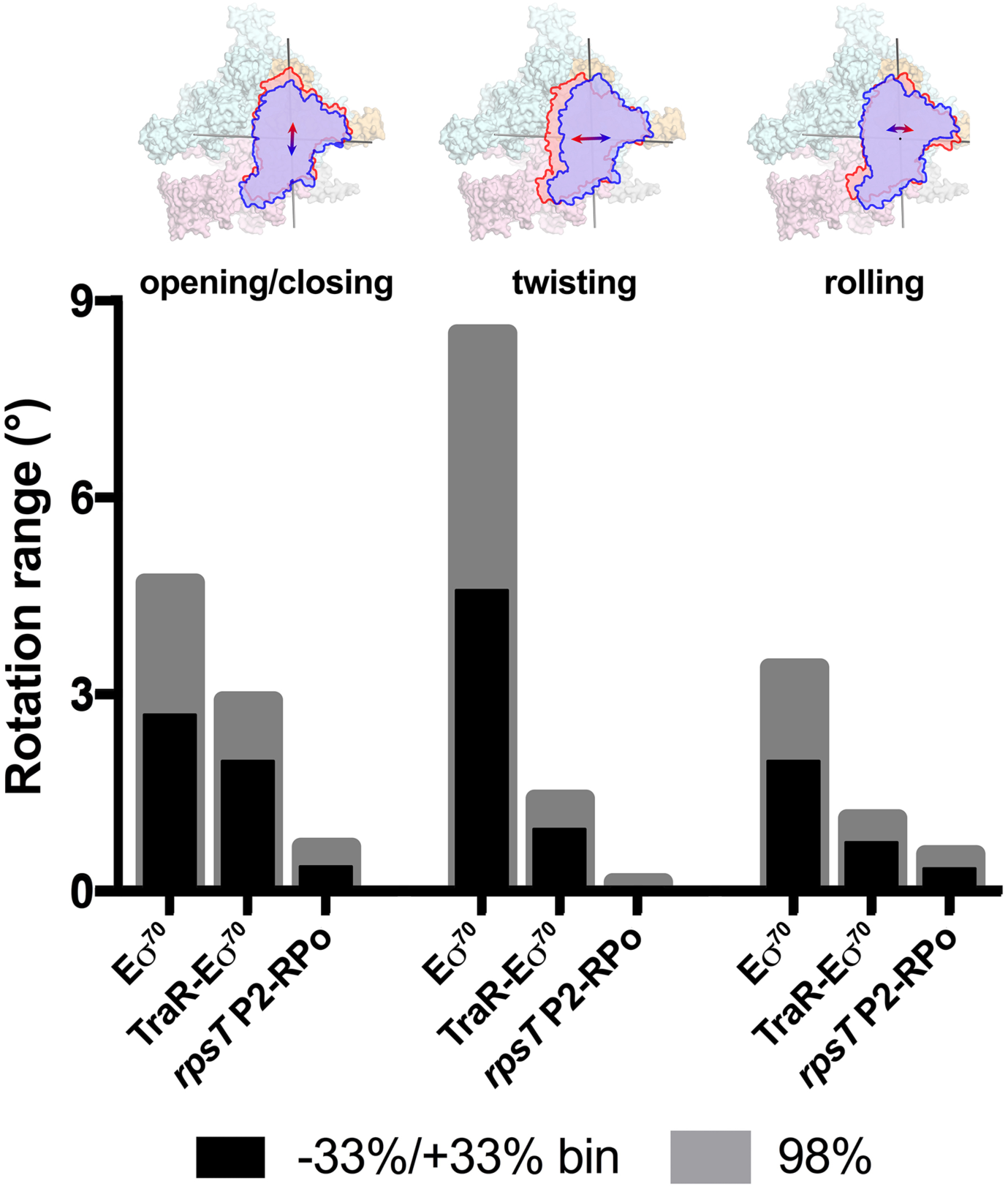
Range of clamp motions for *Eco* RNAP complexes. (top) Eσ^70^ is shown as a molecular surface (α, *ω*, light gray; β, light cyan; β’, light pink; σ^70^, light orange) except the clamp/σ^70^2 module is shown schematically as blue or red outlines (the σ^70^NCR is omitted for clarity) to illustrate the direction and approximate range of motion for the three major components of the clamp motions (left, opening/closing; middle, twisting; left, rolling). (bottom) Histograms denote the range of clamp motions for Eσ^70^, TraR-Eσ^70^, and *rpsT* P2-RPo, as indicated. The black bars denote the range of motion defined by dividing the Eigenvalue histograms into three equal bins and determining the clamp position for the red and blue bins (-33%/+33% bin; see Figure 6). The gray bars denote the estimated range of motion to include 98% of the particles calculated from the Gaussian fits to the Eigenvalue histograms (1% of the particles excluded from each tail; see Figure 6).

These same motions (opening/closing, twisting, rolling) were represented in major components of clamp motion for the TraR-Eσ^70^ and *rpsT* P2-RPo particles as well. The same analyses revealed that TraR binding significantly reduced the range of clamp movement for each of the three clamp motions (Figure 7). We propose that the dampening of clamp motions by TraR facilitates the nucleation of strand opening (Feklistov et al., 2017). As expected, the clamp motions for RPo, with nucleic acids stably bound in the downstream duplex channel, were restricted even further for all three of the major clamp motions (Figure 7).

## Discussion

Our cryo-EM structural analyses show that TraR modulates *Eco* RNAP transcription initiation by binding and altering the conformation and conformational dynamics of the RNAP in four major ways: (1) by manipulating the disposition of β’Si3 (Figures 1A, 1B, 3); (2) by altering the shape of the RNAP active site cleft through a large rearrangement of the βlobe-Si1 (Figure 4); (3) by inducing a significant kink in the BH (Figure 5); and (4) by dampening clamp dynamics (Figures 6, 7; Supplementary movie 1). A previous crystal structure analysis showed that TraR could diffuse into crystals of *Eco* Eσ^70^ and interact with the RNAP β’rim-helices and secondary channel (Molodtsov et al., 2018), but none of these four major TraR-mediated conformational changes seen in the cryo-EM analysis presented here were observed in the crystal structure (Supplementary file 2). Comparing RNAP conformations, the TraR-Eσ^70^ crystal structure (5W1S) matches the Eσ^70^ crystal structure [4YG2, the same crystal form from which the TraR complex was derived; (Murakami, 2013)] much more closely than the TraR-Eσ^70^ cryo-EM structure (Supplementary file 2). Thus, crystal packing forces prevented the conformation of the RNAP from properly responding to TraR binding.

Our results highlight important advantages of cryo-EM over crystallography for structural analysis of large, conformationally dynamic molecular machines such as RNAP (Bai et al., 2015a). First, single-particle cryo-EM analysis does not require crystallization and avoids limitations imposed by crystal packing. Second, multiple, relatively discrete conformational states, such as TraR-Eσ^70^(I), TraR-Eσ^70^(II), and TraR-Eσ^70^(III) (Figures 1A, B, Figure 1 - figure supplement 1), can be revealed from a single sample (Bai et al., 2015b). Third, when a conformational change does not parse into discrete states but rather presents as a continuous distribution of conformations, the range of conformational dynamics can be experimentally assessed (Figures 6, 7) (Nakane et al., 2018).

The consequences of these TraR-induced conformational changes for promoter function (activation or inhibition) depend on the distinctly different properties of the two types of promoters. The kinetics of RPo formation and the thermodynamic properties of RPo vary by many orders of magnitude among different bacterial promoters (McClure, 1985). Eσ^70^ can complete RPo formation on some promoters in a fraction of a second, while other promoters require ten minutes or more. The RPo half-life can vary from a few minutes to many hours. This tremendous range of promoter properties gives rise to a dynamic range for bacterial transcription initiation of ∼4 orders of magnitude and provides rich targets for regulation (Galburt, 2018).

Mechanistic studies of ppGpp/DksA- and TraR-dependent regulation of initiation have revealed general characteristics of promoters that are either activated or inhibited by these factors, and has led to a conceptual model for how TraR activates some promoters while inhibiting others (Gopalkrishnan et al., 2017; Gourse et al., 2018). In the absence of factors, activated promoters generate RPo very slowly (Barker et al., 2001; Paul et al., 2005). Given sufficient time, however, RPo that is ultimately formed is very stable; the activated promoters p*argI*, p*hisG*, and p*thrABC* have half-lives measured in many hours [15 hrs, > 13 hrs, and 6.7 hrs, respectively (Barker et al., 2001)].

On the other hand, inhibited promoters generate RPo very rapidly (Rao et al., 1994). The final transcription-competent RPo is, however, relatively unstable; the RPo half-life of the inhibited promoter *rrnB* P1 is measured in minutes or less (Barker et al., 2001). In the absence of either factors or high initiating NTP concentrations, RPo exists in equilibrium with earlier intermediates along the pathway to RPo formation (Gopalkrishnan et al., 2017; Rutherford et al., 2009).

In order for a transcription factor, such as TraR, to achieve differential regulation (that is, activate some promoters but inhibit others through the same effects on RNAP), the factor must affect more than one feature of the multi-step pathway of RPo formation (Galburt, 2018). In the model for TraR function, TraR acts on all promoters similarly. TraR relieves kinetic barriers to accelerate RPo formation but at the same time likely stabilizes an intermediate prior to RPo formation (Galburt, 2018). Whether TraR activates or inhibits a promoter depends on the basal kinetic landscape for RPo formation of that promoter.

Activated promoters are limited by kinetic barriers that are partially relieved by TraR, accelerating formation of RPo. TraR also likely stabilizes an earlier intermediate relative to RPo, decreasing the RPo half-life. However, the decreased half-life is still measured in hours and so transcription output is limited by other factors and the TraR-mediated acceleration of RPo formation results in activation.

The very short RPo half-life on inhibited promoters means that initiation of RNA chain synthesis competes with dissociation of RPo. High NTP concentrations can shift the equilibrium in favor of RPo by mass action by leading to RNA chain initiation and populating complexes that follow RPo in the transcription cycle (Barker and Gourse, 2001; Murray et al., 2003). Stabilization of an RPo intermediate relative to RPo at these promoters would have a dramatic effect on transcription output by further shifting the occupancy by RNAP to earlier intermediates in the RPo formation pathway (Gopalkrishnan et al., 2017; Rutherford et al., 2009). Although TraR may accelerate RPo formation at these promoters, RPo formation is already fast and transcription output is not limited by this rate. Our structural analysis of the conformational changes imparted on Eσ^70^ by TraR binding and biochemical tests of their functional consequences suggest molecular mechanisms for the effect of TraR on the pathway to RPo formation, providing molecular insight into activation and inhibition.

### Structural mechanism for TraR-mediated activation

While the transcription output of activated promoters is limited by the slow rate of RPo formation, deletion of σ^70^ greatly enhances basal activity (32-fold on p*thrABC*; Figure 4D). This suggests that the presence of σ^70^1.1 in the RNAP channel presents a barrier to the formation of RPo on these promoters. The addition of TraR to EΔ1.1σ^70^ results in a further, small amount of activation, indicating that the presence of σ^70^ is a significant barrier to RPo formation at these promoters but not the only one.

TraR binding induces a large (∼18°) rotation of the βlobe-Si1 module (Figure 4A), distorting the shape of the RNAP channel and altering contacts with σ^70^ in the channel. The TraR-Si1 interaction is essential for weakening σ^70^ contacts, as evidenced by the reduced ability of TraR to activate transcription in the absence of Si1 (Figure 4C). We propose that the TraR interaction with Si1 weakens the σ^70^ -RNAP contacts, facilitating σ^70^ ejection from the channel during RPo formation and thus lowering the kinetic barrier to RPo formation. Our biochemical data indicate that σ^70^ plays a significant role in controlling the overall kinetics at activated promoters but not at inhibited promoters (Figure 4 - figure supplement 1). The contributions of σ^70^ ejection to the overall rates of RPo formation are very different at the two classes of promoters.

Unlike with EΔ1.1σ^70^, TraR together with WT-Eσ^70^ does not result in a 32-fold increase at activated promoters. For example, TraR only activates p*argI* about 3-fold and p*thrABC* about 4-fold under the conditions tested here (Figures 4C, D). This is consistent with our structures where TraR does not cause the complete ejection of σ^70^ from the channel (Figure 1D). Instead, we propose that TraR weakens the σ^70^ -RNAP interactions by shifting the position of the βlobe-Si1 domain, making σ^70^ more easily displaced by the incoming DNA during RPo formation. In other words, this system has evolved to allow modest activation of these promoters with WT-RNAP to a level appropriate for the biological need for these gene products.

Qualitatively, these results are consistent with the interpretation that weakening of the σ^70^ -βlobe interaction allows σ^70^ to be ejected more readily by the incoming DNA during RPo formation, partially relieving the kinetic limitation on activated promoters. Although TraR may also stabilize an intermediate relative to RPo, RPo at these promoters remains relatively stable and RPo formation at these promoters likely results in TraR dissociation.

Our results suggest that the presence of σ^70^ in the DNA channel is a significant obstacle to RPo formation at activated promoters and this step is targeted by TraR, but this is not the only mechanistic determinant of TraR-mediated activation. We found that the formation of the TraR_G_-Si3(II) interface (Figure 3B, C) plays a role in activation, but not inhibition, since deletion of Si3 or mutation of TraR to disrupt the TraR_G_-Si3 interface decreased activation ∼2-fold (compare Figure 3D with 3E) without affecting inhibition. TraR binding also has a significant effect on RNAP clamp dynamics; all three major components of clamp motion in Eσ^70^ were significantly restricted in TraR-Eσ^70^ (Figures 6, 7). We propose that the restriction of clamp motions in TraR-Eσ^70^ could contribute to activation by facilitating transcription bubble nucleation (Feklistov et al., 2017), likely a separate and earlier kinetic step than σ^70^ ejection. The TraR - Si3(SBHMb) interface important for full activation forms in the rotated conformation of Si3 [Si3(II)] in which Si3(SBHMa) contacts with the β’jaw also form. In this way, Si3(II) forms bridging contacts across the RNAP cleft, which may encourage a clamp conformation conducive to more efficient transcription bubble nucleation. Alternatively, the TraR_G_-Si3(SBHMb) interaction may help stabilize the TraR_C_-βlobe-Si1 interaction.

### Structural mechanism for TraR-mediated inhibition

TraR-inhibited promoters have an intrinsically unstable RPo, with earlier intermediates significantly populated at equilibrium (Gopalkrishnan et al., 2017; Rutherford et al., 2009). TraR likely stabilizes one or more of these intermediates relative to RPo, further shifting the equilibrium away from RPo to the intermediate(s) and depopulating RPo. TraR binding induces two distinct conformational changes in the RNAP that we propose disfavor RPo formation, the βlobe-Si1 rotation (Figure 4A) and the BH-kink (Figures 5B, C). Consistent with this hypothesis, ΔSi1-RNAP shows reduced capacity to respond to TraR-mediated effects on inhibition (Figure 4B).

The TraR-mediated βlobe-Si1 rotation (Figure 4A) alters the shape of the RNAP channel, which not only weakens contacts with σ^70^ to help activate positively regulated promoters, but we propose may stabilize DNA contacts in an intermediate prior to RPo. TraR binding also induces a kinked BH which sterically clashes with the proper positioning of the t-strand DNA near the active site (Figure 5). Precise positioning of the t-strand DNA at the active site is critical for efficient catalysis of phosphodiester bond formation by RNAP in the S_N_2 mechanism (Yee et al., 2002). On the basis of TraR-BH contacts observed in the TraR-Eσ^70^ crystal structure, Molodtsov et al. (2018) proposed that TraR-induced BH distortion might affect RPo formation, but other changes inducted by TraR that also contribute to inhibition were not seen in the crystal structure.

Upon the formation of RPo, TraR must dissociate before RNAP can catalyse the first phosphodiester bond because the presence of TraR in the secondary channel sterically blocks NTP binding and TL-folding (Figure 5). We propose that the stable RPo formed at activated promoters is better able to compete wtih TraR binding than the unstable RPo at inhibited promoters (Barker et al., 2001), explaining how TraR-induced BH-kinking could inhibit some promoters but not others.

### TraR manipulates *Eco* RNAP lineage-specific insertions to modulate transcription initiation

The large β and β’ subunits of the bacterial RNAP are conserved throughout evolution, containing 16 and 11 shared sequence regions, respectively, common to all bacterial RNAPs (Lane and Darst, 2010a). These shared sequence regions are separated by relatively nonconserved spacer regions in which large LSIs can occur (Lane and Darst, 2010a). These are typically independently-folded domains, ranging in size from 50 to 500 amino acids, located on the surface of the RNAP and often highly mobile. A key feature of the TraR functional mechanism is modulation of *Eco* RNAP transcription initiation through conformational changes brought about by interactions with two of the *Eco* RNAP LSIs, βSi1 (Figure 4A) and β’Si3 (Figures 3A-C).

Deletions of Eco βSi1 supported basic *in vitro* transcription function and normal *in vivo* cell growth, leading to its original designation as ’dispensable region I’ (Severinov et al., 1994). Later studies revealed that *in vivo*, the ΔβSi1-RNAP was unable to support cell growth at 42°C and could only support slow growth at 30°C (Artsimovitch, 2003). Thus, βSi1 may serve as a binding determinant for unknown transcription regulators that modulate *Eco* RNAP function during unusual growth conditions. Indeed, TraR interacts with Si1 as well as the nearby βlobe to distort the RNAP active site cleft (Figure 4A), effecting both inhibition (Figure 4B) and activation (Figure 4C) by TraR.

Eco β’Si3 is an unusual LSI as it is inserted in the middle of the TL, a key structural element in the RNAP nucleotide addition cycle that is conserved in all multi-subunit RNAPs (Lane and Darst, 2010a). As a consequence, Si3 plays a central role in *Eco* RNAP function and deletion of Si3 is not viable (Artsimovitch, 2003; Zakharova et al., 1998). Si3 is known to be highly mobile, moving to accommodate folding and unfolding of the TL at each RNAP nucleotide addition cycle (Malinen et al., 2012; Zuo and Steitz, 2015); the movement corresponds to a rotation of Si3 by about 33°, resulting in a shift of the Si3 center-of-mass by 15 Å (Kang et al., 2018). Si3 was often disordered in *Eco* RNAP crystal structures [for example, see (Molodtsov et al., 2018)]. In our cryo-EM analysis, TraR engages with Si3, stabilizing a previously unseen conformation of Si3 that plays a role in TraR activation function (Figure 3). Si3 has been implicated previously in RPo formation since the Δβ’Si3-RNAP forms an unstable RPo (Artsimovitch, 2003).

### Conclusion

TraR-like proteins are widespread in proteobacteria and related bacteriophage and plasmids (Gopalkrishnan et al., 2017; Gourse et al., 2018). While the *in vivo* function of TraR is incompletely understood, TraR engages with RNAP in much the same way as DksA/ppGpp, utilizing the same residues in the β’rim-helices that contribute to ppGpp site 2 in the DksA-ppGpp-RNAP complex, and uses its N-terminal α-helix to bind in the RNAP secondary channel near the RNAP active site (Gopalkrishnan et al., 2017; Ross et al., 2016). These general features of TraR binding were confirmed in an X-ray crystal structure of the TraR-Eσ^70^ complex (Molodtsov et al., 2018), but crystal packing constraints prevented this structure from revealing the RNAP conformational changes induced by TraR binding, which are the keys to TraR function. Our structural and functional analyses described here greatly extend previous work (Gopalkrishnan et al., 2017) by identifying the RNAP conformational changes responsible for the effects of TraR on transcription. In so doing, our analysis dissects the complex, multifaceted mechanism that distinguishes activation from inhibition by TraR.

The RPo formation pathway proceeds through multiple steps. TraR binding to RNAP alters the RNAP conformation and conformational dynamics in multiple, complex ways. The complex interplay between TraR binding and RNAP conformation and conformational dynamics allows TraR to modulate multiple features of the energy landscape of RPo formation, which is key to allowing TraR to effect differential regulation across promoter space without direct TraR-promoter interactions.

## Materials & Methods

### Strains, Plasmids and Primer sequences

Plasmids are listed in Supplementary file 3 and oligonucleotide and geneblock sequences are in Supplementary file 4. Bacteria were grown in LB Lennox media or on LB agar plates. Media was supplemented with ampicillin (100 µg/ml) or kanamycin (30 µg/ml) if needed. TraR was made by cloning the *traR* gene in a pET28-based His_10_-SUMO vector which allowed removal of the cleavable N-terminal His_10_-SUMO tag with Ulp1 protease. ESI-Mass Spectrometry revealed that the molecular mass of purified TraR corresponded to that of a monomer lacking the N-terminal methionine [Figure S6 of (Gopalkrishnan et al., 2017)], hence *traR* without the initial M was cloned into the SUMO vector. This tag-less version of TraR exhibited the same level of activity as a previous TraR construct with 4 additional residues (LVPR) at the C-terminal end leftover after His_6_ tag cleavage in the TraR-thrombin site-His_6_ construct (Gopalkrishnan et al., 2017).

### Expression and purification of TraR for cryo-EM

The His_10_-SUMO-TraR plasmid was transformed into competent *Eco* BL21(DE3) by heat shock. The cells were grown in the presence of 25 µg/mL kanamycin to an OD_600_ of 0.5 in a 37°C shaker. TraR expression was induced with a final concentration of 1 mM isopropyl ß-D-thiogalactopyranoside (IPTG) for 3 hours at 37°C. Cells were harvested by centrifugation and resuspended in 50 mM Tris-HCl, pH 8.0, 250 mM NaCl, 5 mM imidazole, 10% glycerol (v/v), 2.5 mM dithiothreitol (DTT), 10 µM ZnCl_2_, 1 mM phenylmethylsulfonyl fluoride (PMSF, Sigma-Aldrich, St. Louis, MO), 1x protease inhibitor cocktail (PIC, Sigma-Aldrich). Cells were homogenized using a continuous-flow French Press (Avestin, Ottawa, ON, Canada) at 4°C and the resulting lysate was centrifuged to isolate the soluble fraction. The supernatant was loaded onto two 5 mL HiTrap IMAC HP columns (GE Healthcare, Pittsburgh, PA) for a total column volume (CV) of 10 mL. His_10_-SUMO-TraR was eluted at 300 mM imidazole in Ni-column buffer [50 mM Tris-HCl, pH 8.0, 500 mM NaCl, 10% glycerol (v/v), 10 µM ZnCl_2_, 2.5 mM DTT]. Peak fractions were combined, treated with ULP1 SUMO-protease overnight, and dialyzed against 20 mM Tris-HCl, pH 8.0, 5% glycerol (v/v), 0.1 mM ethylenediaminetetraacetic acid (EDTA), 500 mM NaCl, 10 µM ZnCl_2_, 2.5 mM DTT, resulting in a final imidazole concentration of 25 mM. The ULP1-cleaved sample was loaded onto one 5 mL HiTrap IMAC HP column to remove His_10_-SUMO-tag along with any remaining uncut TraR. Tagless TraR was collected in the flowthrough and concentrated by centrifugal filtration (Amicon Ultra, EMD Millipore, Burlington, MA). The sample was purified in a final step on a HiLoad 16/60 Superdex 200 column (GE Healthcare). Purified TraR was concentrated to 16 mg/mL by centrifugal filtration, flash-frozen in liquid N_2_, and stored at -80°C.

### *Eco* His10-PPX-RNAP expression and purification

A pET-based plasmid overexpressing each subunit of RNAP (full-length α, β, *ω*) as well as β’-PPX-His_10_ (PPX; PreScission protease site, LEVLFQGP, GE Healthcare) was co-transformed with a pACYCDuet-1 plasmid containing *Eco* rpoZ into *Eco* BL21(DE3). The cells were grown in the presence of 100 µg/mL ampicillin and 34 μg/mL chloramphenicol to an OD_600_ of 0.6 in a 37°C shaker. Protein expression was induced with 1 mM IPTG (final concentration) for 4 hours at 30°C. Cells were harvested by centrifugation and resuspended in 50 mM Tris-HCl, pH 8.0, 5% glycerol (v/v), 10 mM DTT, 1 mM PMSF, and 1x PIC. After French Press lysis at 4°C, the lysate was centrifuged twice for 30 minutes each. Polyethyleneimine [PEI, 10% (w/v), pH 8.0, Acros Organics - ThermFisher Scientific, Waltham, MA] was slowly added to the supernatant to a final concentration of ∼0.6% PEI whith continuous stirring. The mixture was stirred at 4°C for an addition 25 min, then centrifuged for 1.5 hours at 4°C. The pellets were washed three times with 50 mM Tris-HCl, pH 8.0, 500 mM NaCl, 10 mM DTT, 5% glycerol (v/v), 1 mM PMSF, 1x PIC. For each wash, the pellets were homogenized then centrifuged again. RNAP was eluted by washing the pellets three times with 50 mM Tris-HCl, pH 8.0, 1 M NaCl, 10 mM DTT, 5% glycerol (v/v), 1x PIC, 1 mM PMSF. The PEI elutions were combined and precipitated with ammonium sulfate overnight. The mixture was centrifuged and the pellets were resuspended in 20 mM Tris-HCl, pH 8.0, 1 M NaCl, 5% glycerol (v/v), 5 mM DTT. The mixture was loaded onto three 5 mL HiTrap IMAC HP columns for a total CV of 15 ml. RNAP(β’-PPX-His_10_) was eluted at 250 mM imidazole in Ni-column buffer. The eluted RNAP fractions were combined and dialyzed against 20 mM Tris-HCl, pH 8.0, 100 mM NaCl, 5% glycerol (v/v), 5 mM DTT. The sample was then loaded onto a 35 mL Biorex-70 column (Bio-Rad, Hercules, CA), washed with 10 mM Tris-HCl, pH 8.0, 0.1 mM EDTA, 5% glycerol (v/v), 5 mM DTT] in a gradient from 0.2 M to 0.7 M NaCl. The eluted fractions were combined, concentrated by centrifugal filtration, then loaded onto a 320 mL HiLoad 26/600 Superdex 200 column (GE Healthcare) equilibrated in gel filtration buffer [10 mM Tris-HCl, pH 8.0, 0.1 mM EDTA, 0.5 M NaCl, 5% glycerol (v/v), 5 mM DTT]. The eluted RNAP was supplemented with glycerol to 20% (v/v), flash frozen in liquid N_2_, and stored at -80°C.

### *Eco* His10-SUMO-σ^70^ expression and purification

Plasmid encoding *Eco* His_10_-SUMO-σ^70^ was transformed into *Eco* BL21(DE3) by heat shock. The cells were grown in the presence of 50 µg/mL kanamycin to an OD_600_ of 0.6 in 37°C. Protein expression was induced with 1 mM IPTG for 1 hour at 30°C. Cells were harvested by centrifugation and resuspended in 20 mM Tris-HCl, pH 8.0, 5% glycerol (v/v), 500 mM NaCl, 0.1 mM EDTA, 5 mM imidazole, 0.5 mM 2-mercaptoethanol (BME), 1 mM PMSF, 1x PIC. After French Press lysis at 4°C, cell debris was removed by centrifugation. The lysate was loaded onto two 5 mL HiTrap IMAC HP for a total CV of 10 ml. His_10_-SUMO-σ^70^ was eluted at 250 mM imidazole in 20 mM Tris-HCl, pH 8.0, 500 mM NaCl, 0.1 mM EDTA, 5% glycerol (v/v), 0.5 mM BME. Peak fractions were combined, cleaved with ULP1, and dialyzed against 20 mM Tris-HCl pH 8.0, 500 mM NaCl, 0.1 mM EDTA, 5% glycerol (v/v), 0.5 mM BME, resulting in a final imidazole concentration of 25 mM. The cleaved sample was loaded onto one 5 mL HiTrap IMAC HP to remove His_10_-SUMO-tag along with any remaining uncut σ^70^. Tagless σ^70^ was collected in the flowthrough and concentrated by centrifugal filtration. The sample was then loaded onto a HiLoad 16/60 Superdex 200 in gel filtration buffer. Peak fractions of σ^70^ were pooled, supplemented with glycerol to a final concentration of 20% (v/v), flash-frozen in liquid N_2_, and stored at -80°C.

### Preparation of Eσ^70^ for cryo-EM

Eσ^70^ was formed by mixing purified RNAP and 2.5-fold molar excess of σ^70^ and incubating for 20 minutes at 37°C. Eσ^70^ was purified on a Superose 6 Increase 10/300 GL column (GE Healthcare) in gel filtration buffer (10 mM Tris-HCl, pH 8.0, 200 mM KCl, 5 mM MgCl_2_, 10 µM ZnCl_2_, 2.5 mM DTT). The eluted Eσ^70^ was concentrated to ∼10 mg/mL (∼21 μM) by centrifugal filtration (Amicon Ultra).

### Preparation of TraR-Eσ^70^ for cryo-EM

Eσ^70^ was formed by mixing purified RNAP and a 2-fold molar excess of σ^70^ and incubating for 15 minutes at room temperature. Eσ^70^ was purified over a Superose 6 Increase 10/300 GL column in gel filtration buffer. The eluted Eσ^70^ was concentrated to ∼5.0 mg/mL (∼10 μM) by centrifugal filtration. Purified TraR was added (5-fold molar excess over RNAP) and the sample was incubated for 15 min at room temperature. An *rrnB* P1 promoter fragment (Integrated DNA Technologies, Coralville, IA) was was added (2-fold molar excess over RNAP) and the sample was incubated for a further 15 minutes at room temperature. The *rrnB* P1 promoter fragment did not bind to TraR-Eσ^70^ under the cryo-EM grid preparation conditions - the subsequent structural analyses did not reveal any evidence of promoter binding.

### Preparation of *rpsT* P2-RPo for cryo-EM

Eσ^70^ was prepared as described for TraR-Eσ^70^, but after the size exclusion chromatography the complex was concentrated to ∼10 mg/mL (∼20 μM) by centrifugal filtration. Duplex *rpsT* P2 promoter fragment (-60 to +25, Figure 3A, IDT) was added to the concentrated Eσ^70^ to 3-fold molar excess. The sample was incubated for 20 mins at room temperature prior to cryo-EM grid preparation.

### Cryo-EM grid preparation

CHAPSO {3-([3-cholamidopropyl]dimethylammonio)-2-hydroxy-1-propanesulfonate} (Anatrace, Maumee, OH) was added to the samples to a final concentration of 8 mM (Chen et al., 2019). The final buffer condition for all the cryo-EM samples was 10 mM Tris-HCl, pH 8.0, 100 mM KCl, 5 mM MgCl_2_, 10 μM ZnCl_2_, 2.5 mM DTT, 8 mM CHAPSO. C-flat holey carbon grids (CF-1.2/1.3-4Au) were glow-discharged for 20 sec prior to the application of 3.5 μL of the samples. Using a Vitrobot Mark IV (FEI, Hillsboro, OR), grids were blotted and plunge-froze into liquid ethane with 100% chamber humidity at 22°C.

### Acquisition and processing of TraR-Eσ^70^ cryo-EM dataset

Grids were imaged using a 300 keV Krios (FEI) equipped with a K2 Summit direct electron detector (Gatan, Pleasanton, CA). Datasets were recorded with Serial EM (Mastronarde, 2005) with a pixel size of 1.3 Å over a defocus range of 0.8 μm to 2.4 μm. Movies were recorded in counting mode at 8 electrons/physical pixel/second in dose-fractionation mode with subframes of 0.3 sec over a 15 sec exposure (50 frames) to give a total dose of 120 electrons/physical pixel. Dose-fractionated movies were gain-normalized, drifted-corrected, summed, and dose-weighted using MotionCor2 (Grant and Grigorieff, 2015; Zheng et al., 2017). CTFFIND4 (Rohou and Grigorieff, 2015) was used for contrast transfer function estimation. Particles were picked using Gautomatch (http://www.mrc-lmb.cam.ac.uk/kzhang/) using a 2D template. Picked particles were extracted from the dose-weighted images with RELION (Zivanov et al., 2018) using a box size of 256 pixels. Two TraR-Eσ^70^ datasets were collected: dataset 1 consisted of 1,546 motion-corrected images with 631,880 particles and dataset 2 consisted of 2,132 motion-corrected images with 378,987 particles. The particles from each dataset were curated using RELION 3D classification (N=3) using a cryoSPARC ab-initio reconstruction (Punjani et al., 2017) generated from a subset of the particles. The highest resolution classes from each dataset were subjected to RELION 3D auto-refinement resulting in a 4.69 Å resolution map from dataset 1 and a 4.38 Å resolution map from dataset 2. Refinement metadata and post-processing were used as inputs for RELION CTF refinement and Bayesian Polishing (Zivanov et al., 2018). The polished particles from both datasets were combined, resulting in 372,670 particles. The particles were aligned using RELION 3D auto-refinement resulting in a consensus map with nominal resolution of 3.62 Å. Using the refinement parameters, subtractive 3D classification (N=3) was performed on the particles by subtracting density outside of β’Si3 and classifying in a mask around β’Si3. Classification revealed three distinct β’Si3 dispositions (Figure S1D). Local refinement metadata (highlighted in red dotted box, Figure S1D) for TraR-Eσ^70^(I) and TraR-Eσ^70^(II) were used for RELION multi-body refinements to examine clamp motions (Nakane et al., 2018). Local resolution calculations were performed using blocres and blocfilt from the Bsoft package (Cardone et al., 2013).

### Acquisition and processing of Eσ^70^ cryo-EM dataset

The Eσ^70^ image acquisition and processing was the same as for TraR-Eσ^70^ except with the following differences. Grids were imaged using a 200 keV Talos Arctica (FEI) equipped with a K2 Summit direct electron detector. Datasets were recorded with a pixel size of 1.3 Å over a defocus range of -1.0 μm to -2.5 μm. Movies were recorded in counting mode at 8.4 electrons/physical pixel/second in dose-fractionation mode with subframes of 0.2 sec over a 10 sec exposure (50 frames) to give a total dose of 84 electrons/physical pixel. Picked particles were extracted from the dose-weighted images in RELION (Scheres 2012) using a box size of 200 pixels. The Eσ^70^ dataset consisted of 3,548 motion-corrected images with 1,387,166 particles. A subset of the particles was subjected to cryoSPARC ab-initio reconstruction (Punjani 2017) to generate a 3D template for 3D classifications in cryoSPARC and 3D refinements in RELION (Scheres 2012). Particles were split into two groups (1^st^ group: particles from images 1-2,000; 2^nd^ group: particles from images 2001-3548. Particles from each group were curated using cryoSPARC heterogeneous refinement (N=3) resulting in a subset of 479,601 particles for the first group and 329,293 particles for the second group. Curated particles were combined and a consensus refinement was performed in RELION using the cryoSPARC generated initial model resulting in a map with nominal resolution of 4.54 Å (without post-processing). Particles from this refinement (highlighted in red dotted box, Figure 1 - figure supplement 3) were further analyzed using RELION multi-body refinement as described in the text (Nakane et al., 2018). Additionally, particles were further curated using RELION 3D classification (N=3) without alignment. Classification revealed two lower resolution class and a higher resolution class. The higher resolution class containing 358,725 particles was RELION 3D auto-refined and subjected to RELION CTF refinement and RELION Bayesian Polishing (Zivanov et al., 2018). After polishing, particles were refined to a nominal resolution of 4.05 Å after RELION post-processing.

### Acquisition and processing of *rpsT* P2-RPo cryo-EM dataset

The *rpsT* P2-RPo cryo-EM image acquisition and processing was the same as for TraR-Eσ^70^ except with the following differences. The imaging defocus range was 0.5 μm to 2.5 μm. Movies were recorded in super-resolution mode at 8 electrons/physical pixel/second in dose-fractionation mode with subframes of 0.2 sec over a 10 sec exposure (50 frames) to give a total dose of 80 electrons/physical pixel. The *rpsT* P2-RPo dataset consisted of 6,912 motion-corrected images with 973,481 particles. In RELION, a consensus refinement was performed using the extracted particles and a cryoSPARC generated initial model resulting in a 4.62 Å resolution map. Using the refinement parameters, 3D classification (N=2) was performed on the particles without alignment. Classification revealed a lower resolution class and a higher resolution class of with 370,965 particles with nominal resolution of 4.38 Å after RELION 3D auto-refinement. Refinement metadata and post-processing were used as inputs for RELION CTF refinement and RELION Bayesian Polishing (Zivanov et al., 2018). Subsequent 3D classification (N=3) was used to further classify the polished particles resulting in one junk class and two high resolution classes (Figure 2 - figure supplement 1). The highest resolution reconstruction (3.43 Å) contained 289,679 particles.

### Model building and refinement of cryo-EM structures

To build initial models of the protein components of the complexes, a crystal structure of *Eco* Eσ^70^ [PDB ID 4LJZ, with σ^70^ from 4LK1; (Bae et al., 2013)] was manually fit into the cryo-EM density maps using Chimera (Pettersen et al., 2004) and manually adjusted using Coot (Emsley and Cowtan, 2004). For TraR-Eσ^70^, σ^70^1.1 from 4LK1 (Bae et al., 2013) and TraR from 5W1S (Molodtsov et al., 2018) were also added. For *rpsT* P2-RPo, the promoter DNA was manually added. Appropriate domains of each complex were rigid-body refined, then subsequently refined with secondary structure and nucleic acid restraints using PHENIX real space refinement (Adams et al., 2010).

### Purification of TraR and RNAP for transcription assays

IPTG (1 mM final) was used to induce expression of TraR (WT or variant) from *Eco* BL21 DE3 *dksA*::Tn10 (RLG7075) host cells. TraR and variants were purified as described (Gopalkrishnan et al., 2017), either from His_6_-TraR overexpression plasmids with removal of the His_6_-tag with thrombin, or from His_10_-SUMO-TraR constructs with removal of the His_10_-SUMO-tag with Ulp1 protease, resulting in a 72 amino acid TraR lacking the N-terminal Met. WT-TraR purified by the two methods gave comparable results. WT and variant RNAPs were purified as described previously (Ross et al., 2016). The Δ1.1σ^70^ was expressed and purified as described previously (Chen et al., 2017). EΔ1.1σ^70^ was reconstituted with a 4:1 molar ratio of Δ1.1σ^70^ to core RNAP. The purified core RNAP lacked detectable WT-σ^70^ activity.

### *In Vitro* transcription assays, site-directed mutagenesis, and TraR-RNAP binding assays

All of these procedures were carried out exactly as previously described (Gopalkrishnan et al., 2017).

## Acknowledgments

We thank M. Ebrahim and J. Sotiris at The Rockefeller University Evelyn Gruss Lipper Cryo-electron Microscopy Resource Center for help with cryo-EM data collection, and R. Saecker and other members of the Darst-Campbell Laboratory for helpful discussions. This work was supported by NIH grants R01 GM114450 to E.A.C., R01 GM37048 to R.L.G. and R35 GM118130 to S.A.D.

## Accession Numbers

The cryo-EM density maps have been deposited in the EMDataBank under accession codes EMD-0348 [*Eco* TraR-Eσ^70^(I)], EMD-0349 [*Eco* TraR-Eσ^70^(II)], EMD-20231 [*Eco* TraR-Eσ^70^(III)], EMD-20230 (*Eco* Eσ^70^), EMD-20203 (*rpsT* P2-RPo), and EMD-20232 (*rpsT* P2-RPo2). The atomic coordinates have been deposited in the Protein Data Bank under accession codes 6N57 [*Eco* TraR-Eσ^70^(I)], 6N58 [*Eco* TraR-Eσ^70^(II)], 6P1K (*Eco* Eσ^70^), and 6OUL (*rpsT* P2-RPo).

---

**Figure 1 - figure supplement 1.**
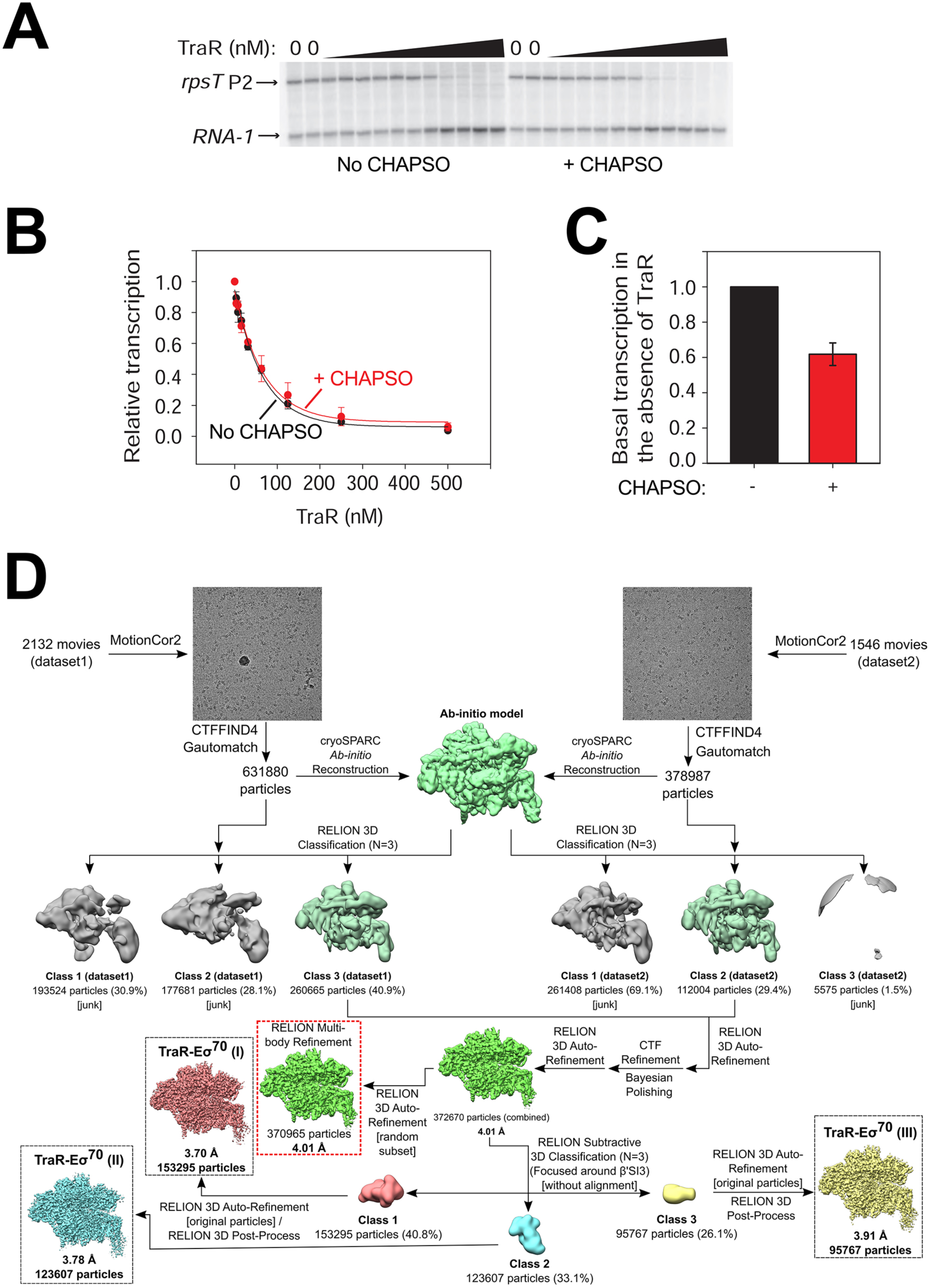
Cryo-EM solution conditions do not affect TraR function and TraR-Eσ^70^ cryo-EM processing pipeline. **(A)** Multi-round *in vitro* transcription of *rpsT* P2 by Eσ^70^ (20 nM) at a range of TraR concentrations (wedge indicates 4 nM - 4 µM) in the absence or presence of 8 mM CHAPSO as indicated. Plasmid templates also contained the *RNA-1* promoter **(B)** Quantification of transcripts from experiments like those in **(A)** plotted relative to values in the absence of TraR. The IC_50_ for inhibition by TraR was ∼50 nM for both ± CHAPSO data sets. Averages with range from two independent experiments are plotted. **(C)** Transcription in the absence of TraR is plotted, relative to the same reactions without CHAPSO. Although it had no effect on the concentration of TraR required for half-maximal inhibition (Figure S1B), CHAPSO reduced transcription slightly. Averages with range from two independent experiments are plotted. **(D)** TraR-Eσ^70^ cryo-EM processing pipeline.

**Figure 1 - figure supplement 2.**
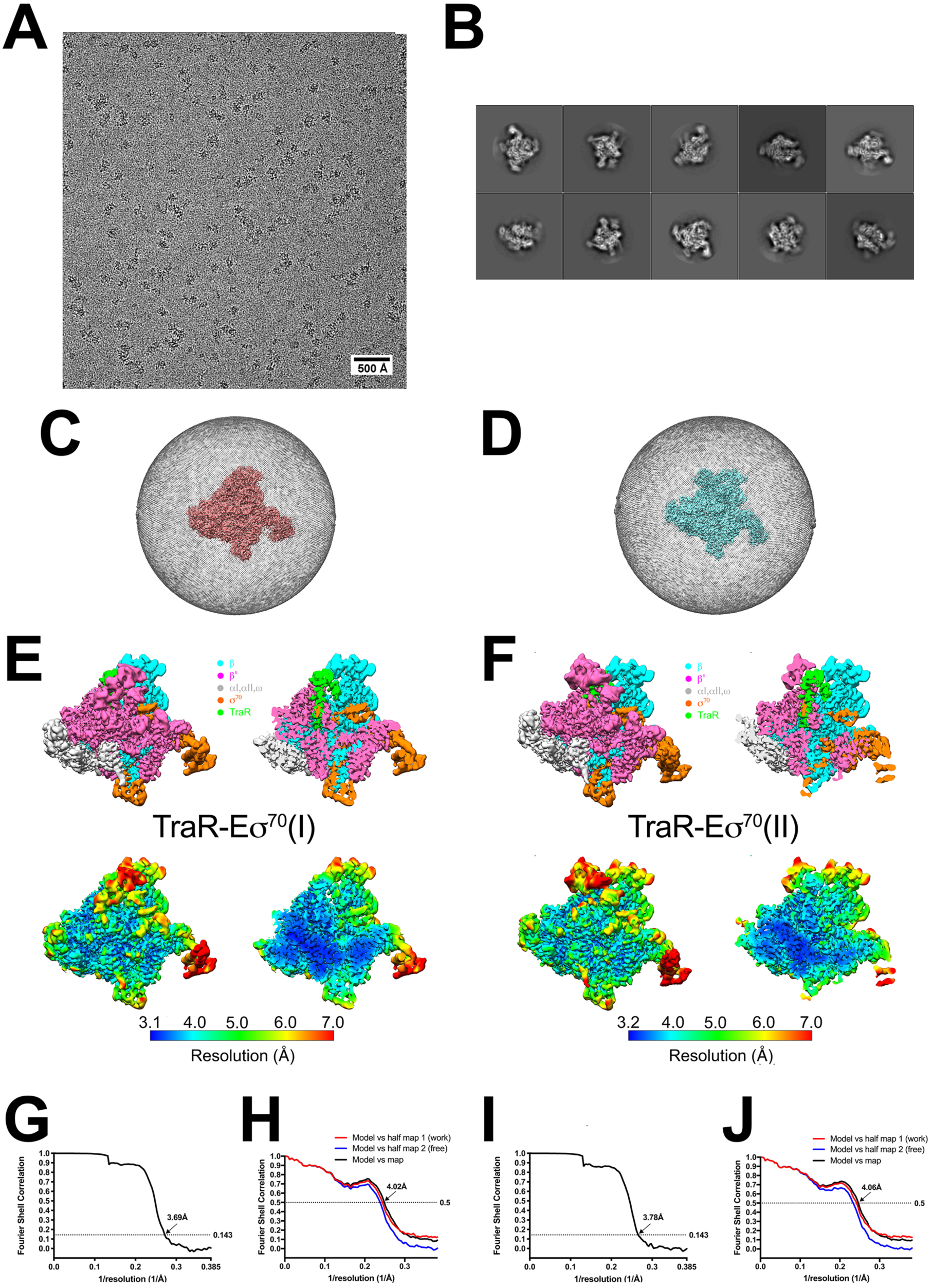
TraR-Eσ^70^ cryo-EM. A. Representative micrograph of TraR-Eσ^70^ in vitreous ice. B. The ten most populated classes from 2D classification. C. Angular distribution for TraR-Eσ^70^(I) particle projections. D. Angular distribution for TraR-Eσ^70^(II) particle projections. E. (top) The 3.7-Å resolution cryo-EM density map of TraR-Eσ^70^(I) is colored according to the key. The right view is a cross-section of the left view. (bottom) Same views as (top) but colored by local resolution (Cardone et al., 2013). F. (top) The 3.8-Å resolution cryo-EM density map of TraR-Eσ^70^(II) is colored according to the key. The right view is a cross-section of the left view. (bottom) Same views as (top) but colored by local resolution (Cardone et al., 2013). G. Gold-standard FSC of TraR-Eσ^70^(I). The gold-standard FSC was calculated by comparing the two independently determined half-maps from RELION. The dotted line represents the 0.143 FSC cutoff, which indicates a nominal resolution of 3.7 Å. H. FSC calculated between the refined structure and the half map used for refinement (work), the other half map (free), and the full map. I. Gold-standard FSC of TraR-Eσ^70^(II). The gold-standard FSC was calculated by comparing the two independently determined half-maps from RELION. The dotted line represents the 0.143 FSC cutoff, which indicates a nominal resolution of 3.8 Å. J. FSC calculated between the refined structure and the half map used for refinement (work), the other half map (free), and the full map.

**Figure 1 - figure supplement 3.**
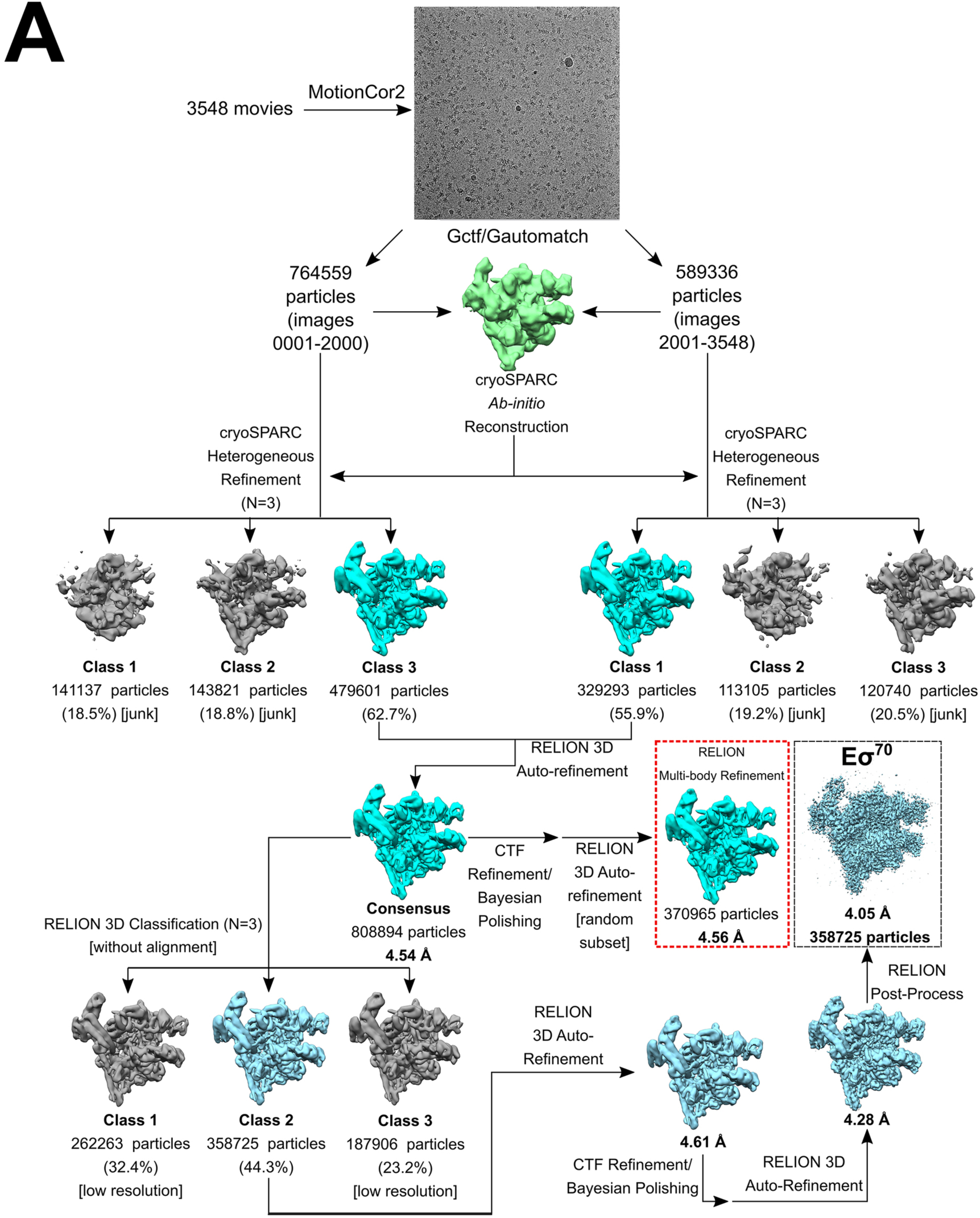
Eσ^70^ cryo-EM processing pipeline. Eσ^70^ cryo-EM processing pipeline.

**Figure 1 - figure supplement 4.**
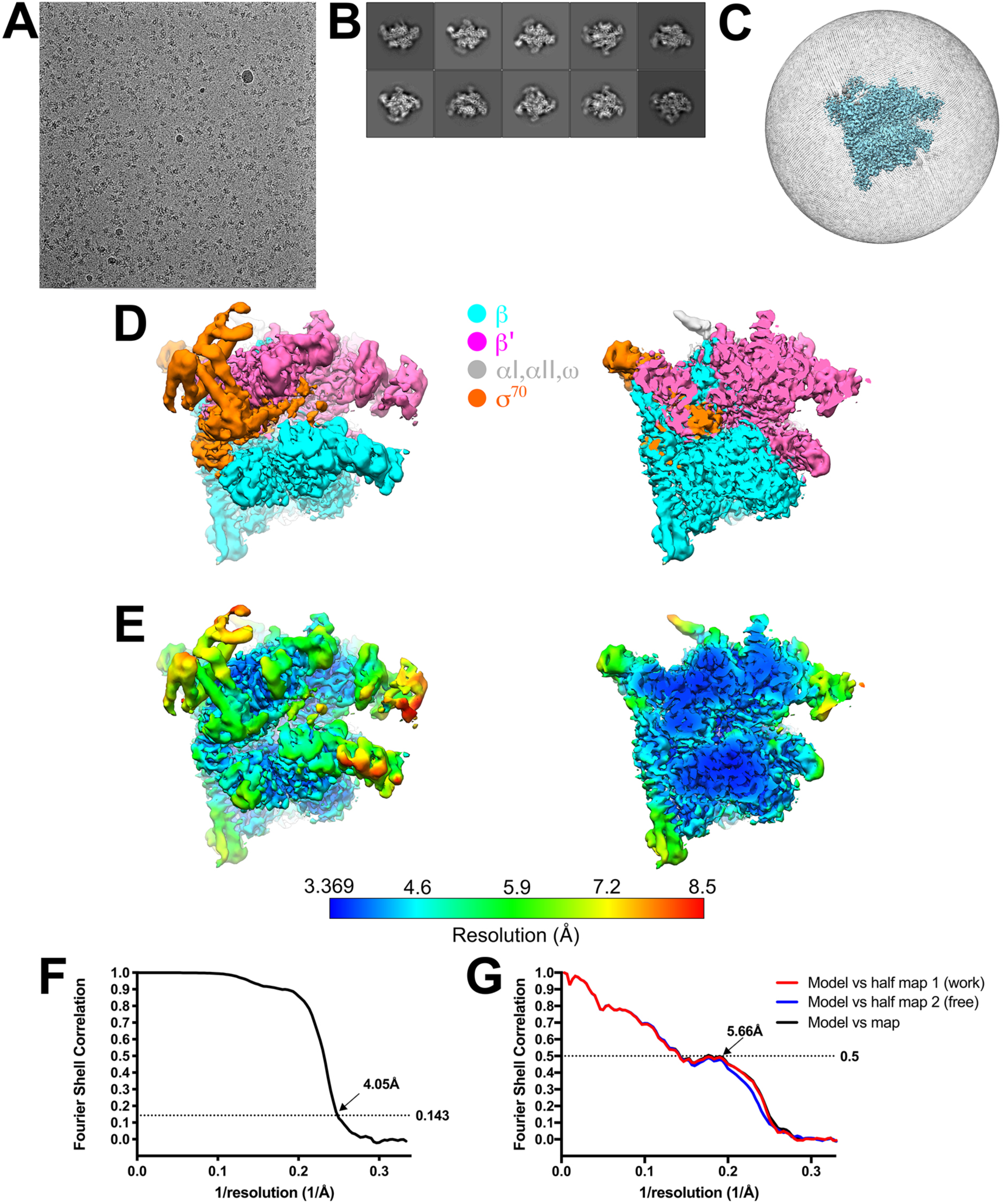
Eσ^70^ cryo-EM. A. Representative micrograph of Eσ^70^ in vitreous ice. B. The ten most populated classes from 2D classification. C. Angular distribution for Eσ^70^ particle projections. D. The 4.1-Å resolution cryo-EM density map of Eσ^70^ is colored according to the key. The right view is a cross-section of the left view. E. Same views as (D) but colored by local resolution (Cardone et al., 2013). F. Gold-standard FSC of Eσ^70^. The gold-standard FSC was calculated by comparing the two independently determined half-maps from RELION. The dotted line represents the 0.143 FSC cutoff, which indicates a nominal resolution of 4.1 Å. G. FSC calculated between the refined structure and the half map used for refinement (work), the other half map (free), and the full map.

**Figure 2 - figure supplement 1.**
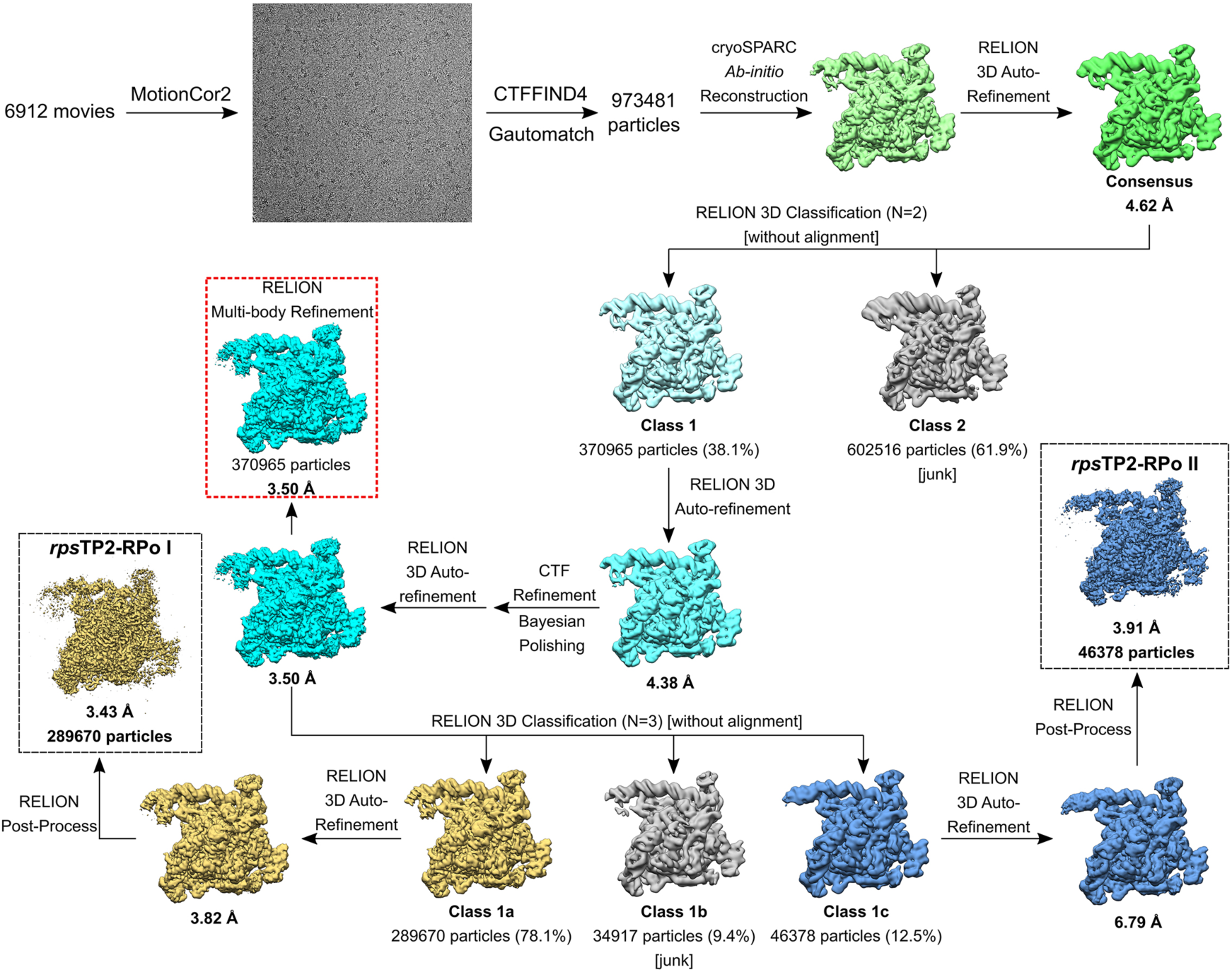
*rpsT* P2-RPo cryo-EM processing pipeline.

**Figure 2 - figure supplement 2.**
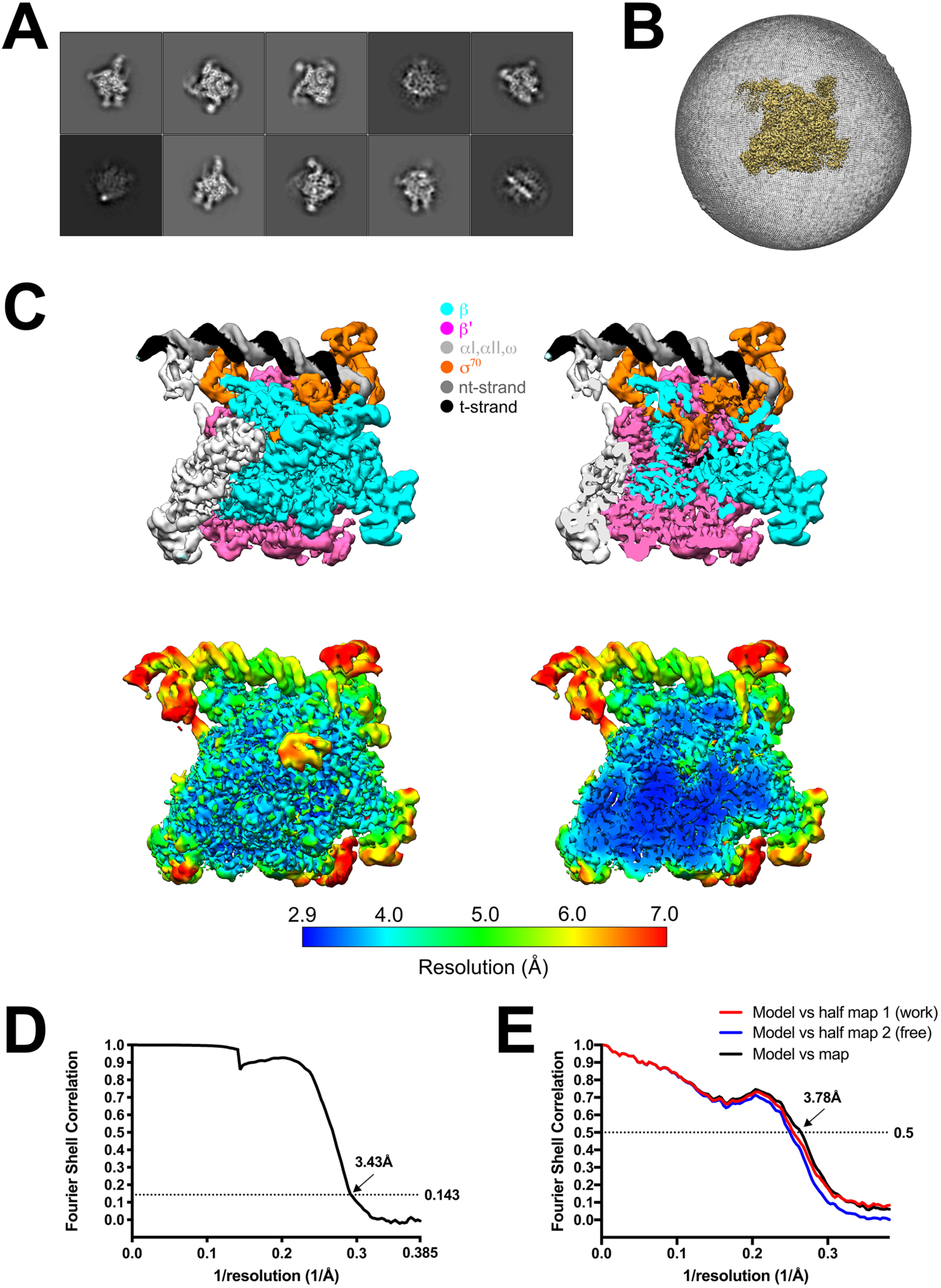
*rpsT* P2-RPo cryo-EM. A. The ten most populated classes from 2D classification. B. Angular distribution for *rpsT* P2-RPo particle projections. C. (top) The 3.4-Å resolution cryo-EM density map of *rpsT* P2-RPo is colored according to the key. The right view is a cross-section of the left view. (bottom) Same views as (top) but colored by local resolution (Cardone et al., 2013). D. Gold-standard FSC of *rpsT* P2-RPo. The gold-standard FSC was calculated by comparing the two independently determined half-maps from RELION. The dotted line represents the 0.143 FSC cutoff, which indicates a nominal resolution of 3.4 Å. E. FSC calculated between the refined structure and the half map used for refinement (work), the other half map (free), and the full map.

**Figure 3 - figure supplement 1.**
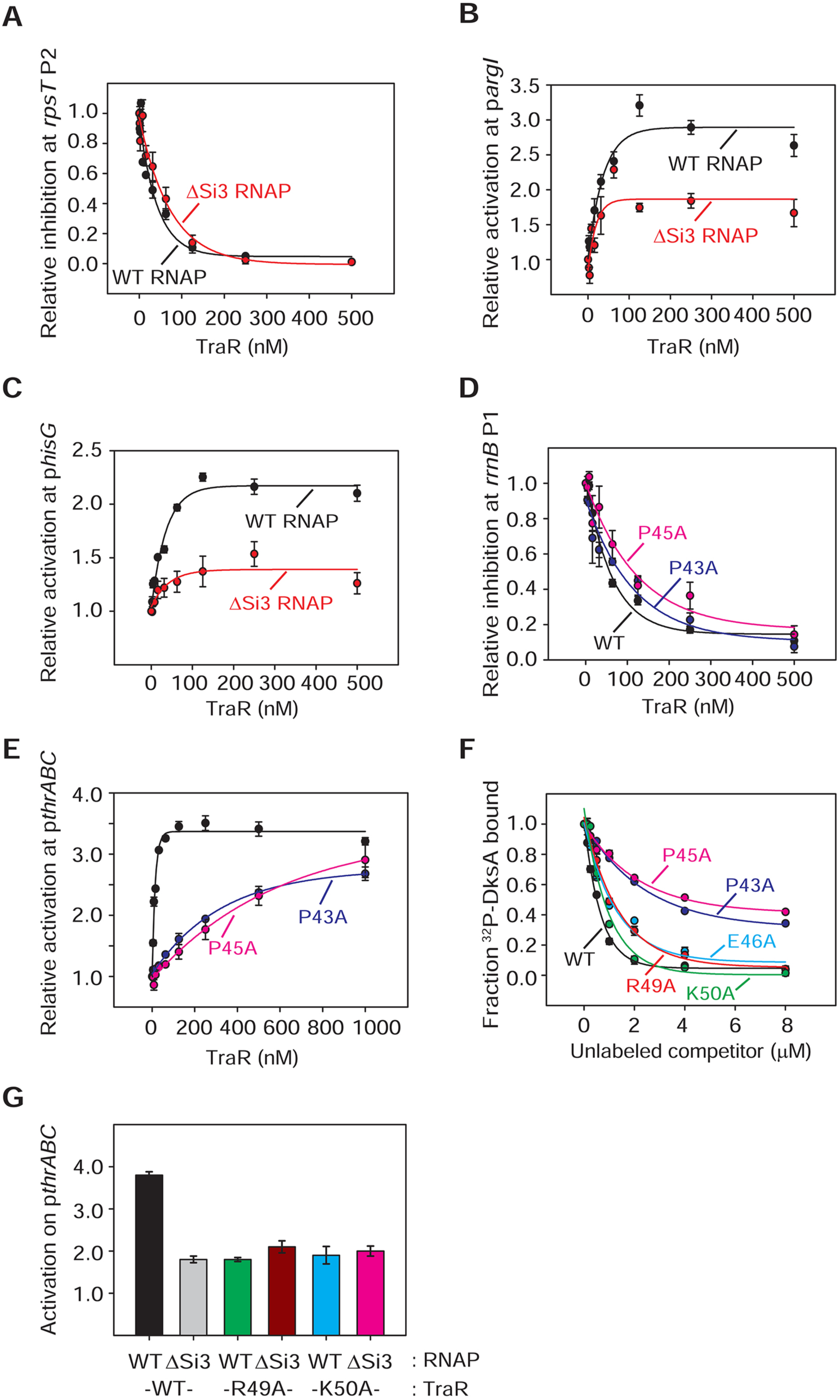
RNAP-Si3 interaction with TraR_G_ residues. (**A**) - (**G**) Quantifications show averages with range from two independent experiments. **(A) (B) (C)** Multi round *in vitro* transcription of *rpsT* P2 (**A**), p*argI* (**B**) or p*hisG* (**C**) was performed at a range of TraR concentrations (2 nM - 2 µM) in the presence of 20 nM WT- or ΔSi3-RNAP. Transcripts were quantified and plotted relative to values in the absence of TraR. Averages with range from two independent experiments are shown. **(A)** The IC_50_ for inhibition by TraR was ∼60 nM with WT-RNAP and ∼90 nM with ΔSi3-RNAP. **(D) (E)** Multi round *in vitro* transcription from *rrnB* P1 **(D)** and p*thrABC* **(E)** was performed with 20 nM WT-Eσ^70^ at a range of concentrations of WT- or variant TraR (2 nM - 2 µM). Transcripts were quantified and plotted relative to values in the absence of TraR. Error bars denote the standard deviation of three independent measurements. For (**D**), the IC_50_ for inhibition by WT-TraR was ∼50 nM, by P43A-TraR was ∼80 nM, and by P45A-TraR was ∼115 nM. **(F)** Effect of substitutions in TraR_G_ residues on binding to RNAP was determined by competition with ^32^P-DksA in an Fe^2+^-mediated cleavage assay. WT-TraR (∼0.6 µM), P43A-TraR (∼3 μM), P45A-TraR (∼4 μM), E46A-TraR (∼1 μM), R49A-TraR (∼1 μM) and K50A-TraR (∼0.7 μM) reduced cleavage of 1.0 mM ^32^P-DksA by 50%. Averages with range from two independent experiments are shown. **(G)** Transcription experiments were carried out with 20 nM WT- or ΔSi3-RNAP with 250 nM WT- or variant TraR as indicated. Values are relative to basal transcription by WT-RNAP without factor (normalized to1.0). Averages with range from two independent experiments are shown.

**Figure 4 - figure supplement 1.**
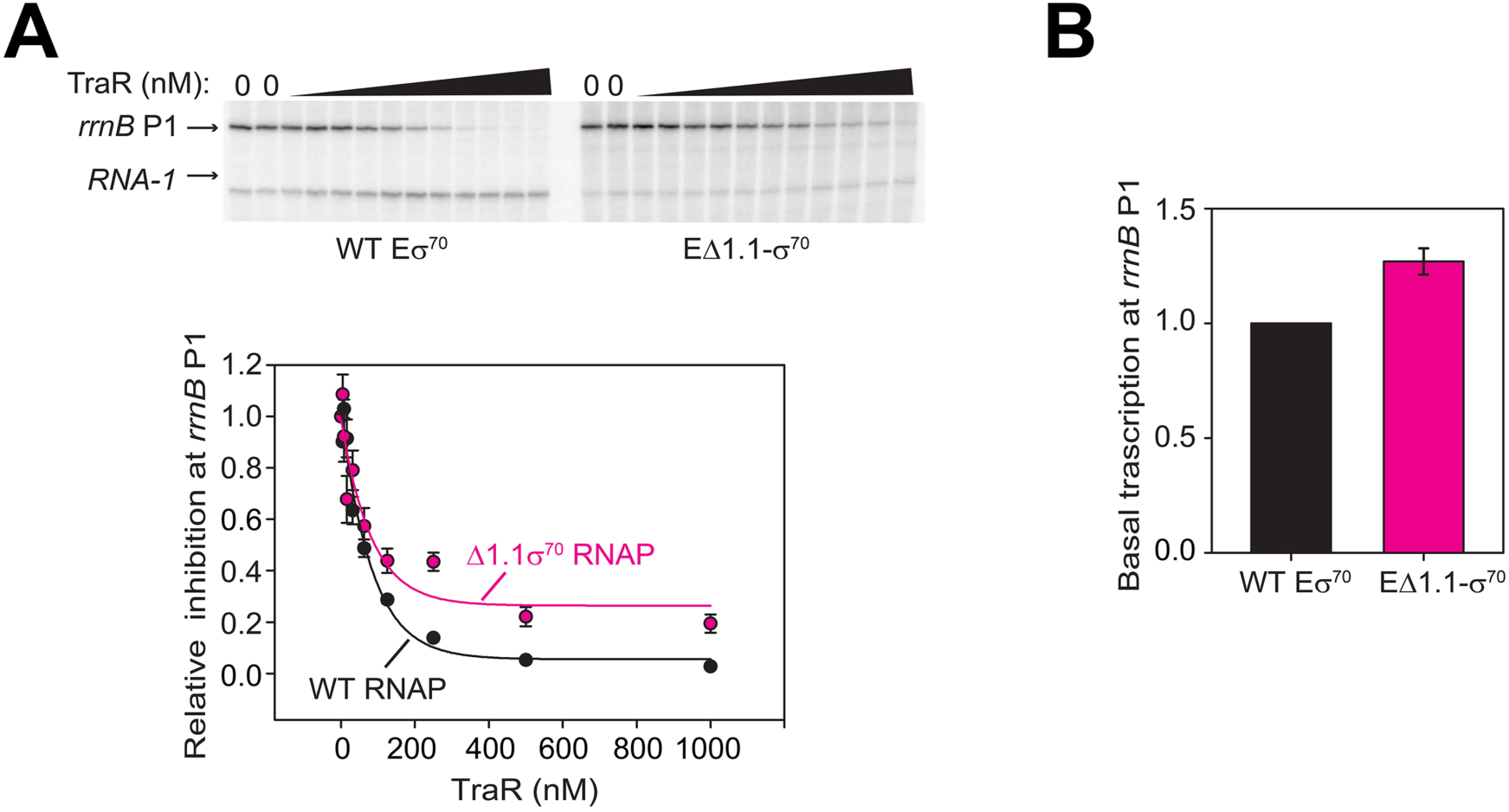
EΔ1.1σ^70^ has small defects for inhibition of *rrnB* P1 by TraR. **(A)** (top) Multi round *in vitro* transcription at *rrnB* P1 was carried out at a range of TraR concentrations (wedge indicates 4 nM - 4 µM) in the presence of 20 nM WT-Eσ^70^ or EΔ1.1σ^70^ as indicated. Plasmid templates also contained the *RNA-1* promoter. (bottom) Transcripts were quantified and plotted relative to values in the absence of TraR. The IC_50_ for inhibition by TraR was ∼50 nM with WT-RNAP and ∼90 nM with Δ1.1σ^70^-RNAP. Averages with range from two independent experiments are shown. **(B)** Basal level of transcription from *rrnB* P1 is only slightly affected by Δ1.1σ^70^. Error bars denote standard deviation of three independent measurements.

**Supplementary file 1.**
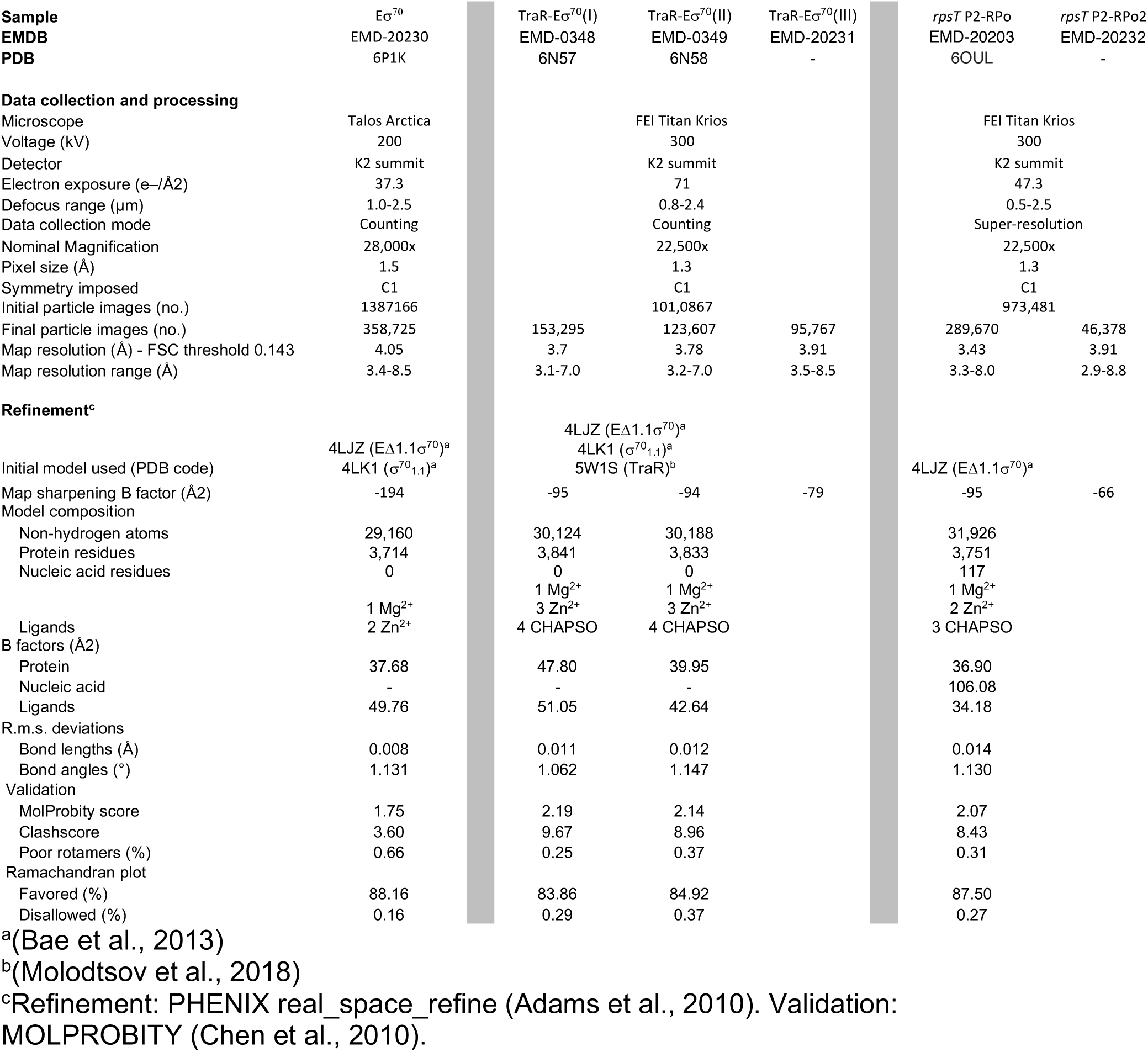
Cryo-EM data collection and refinement.

**Supplementary file 2.**
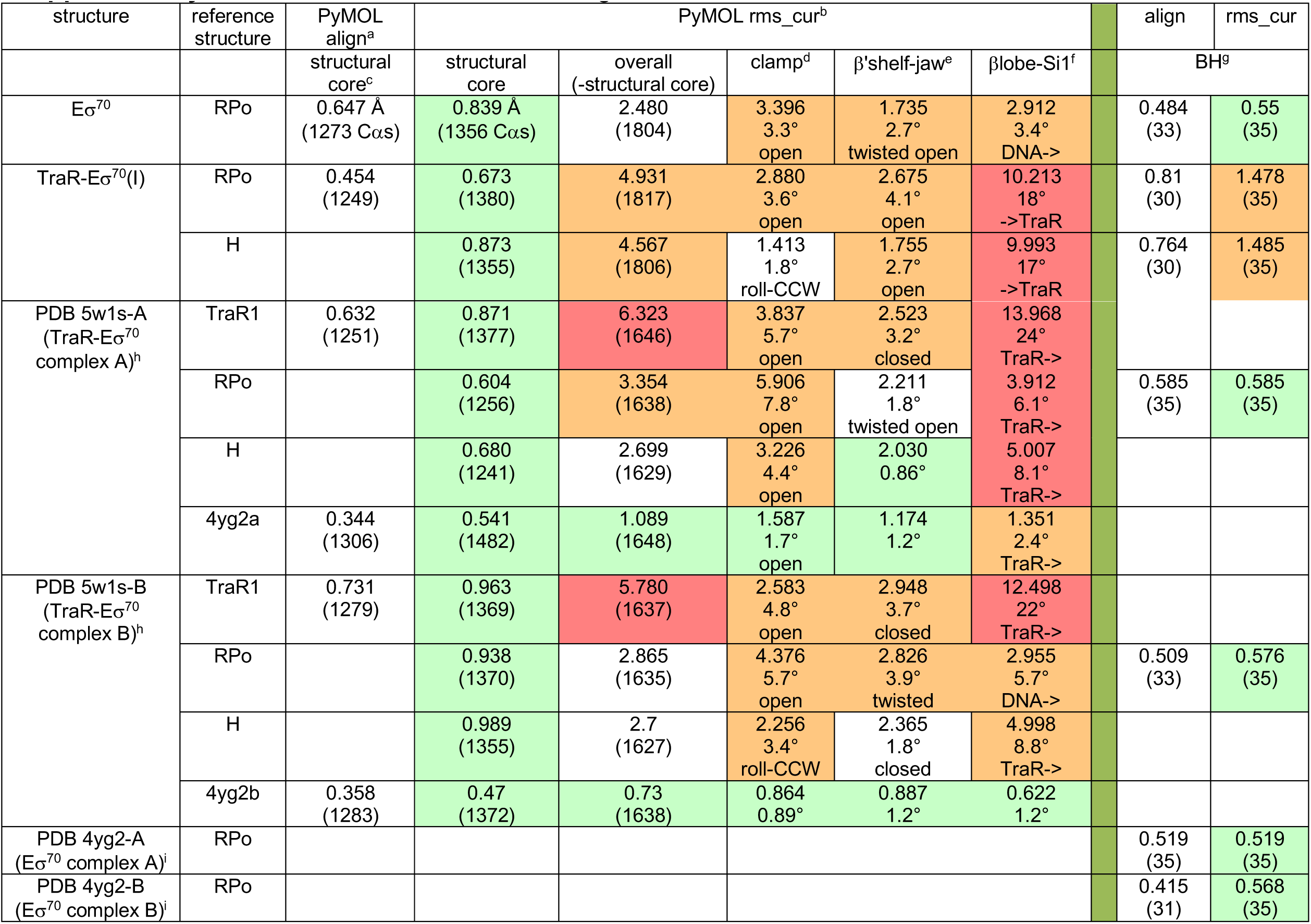
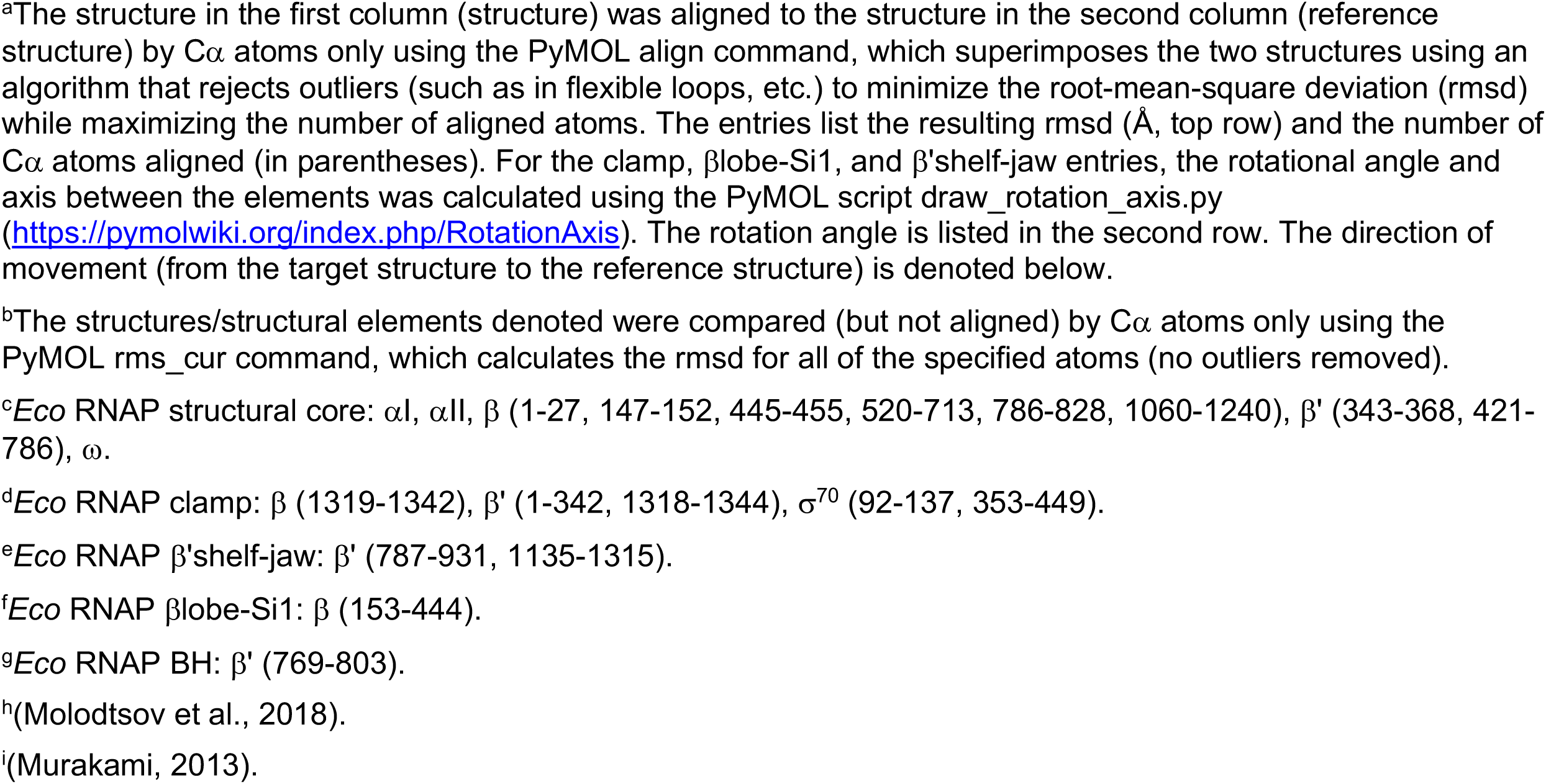
RNAP conformational changes.

**Supplementary file 3.**
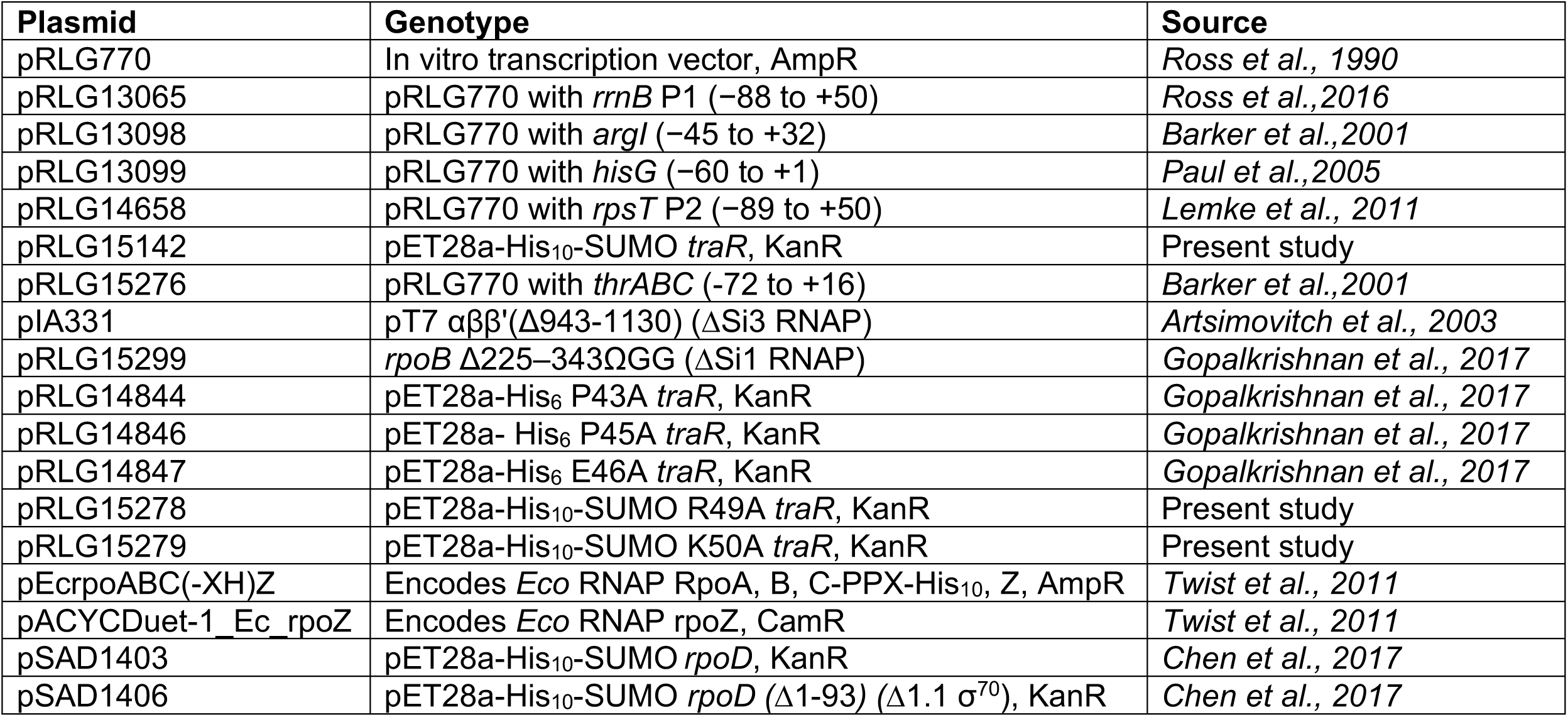
Plasmids used in this study.

**Supplementary file 4.**
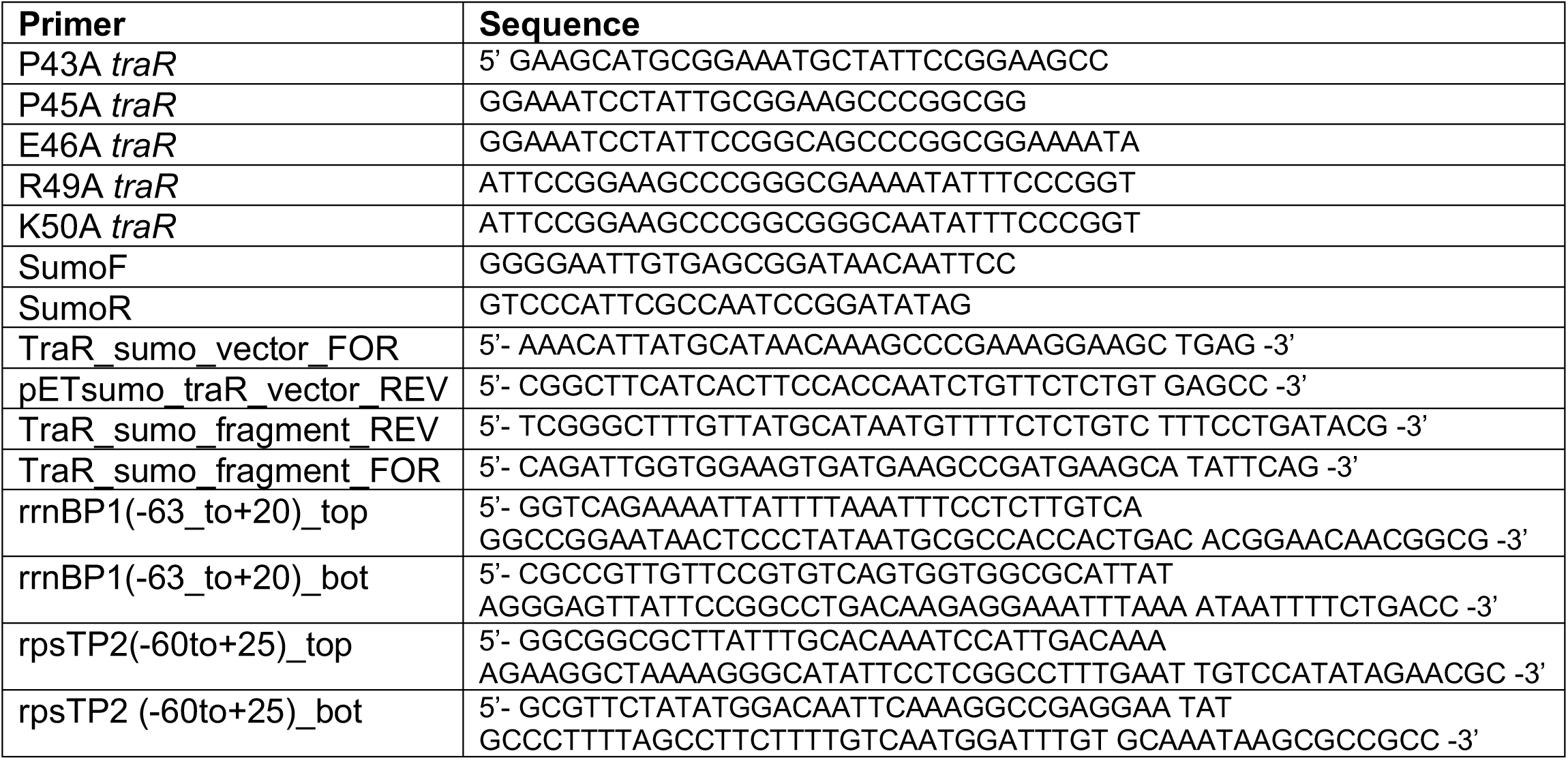
Plasmids used in this study.

**Supplementary movie 1.** Movie illustrating changes in conformation and conformational dynamics of RNAP induced by TraR binding.

